# The cost-benefit trade-off of peritrichous flagellation in bacteria

**DOI:** 10.64898/2026.08.20.746045

**Authors:** María José Giralt-Zúñiga, Michael Jahn, Joshua L. Franklin, Kathirvel Alagesan, Florian Kondrot, Eugen Kaganovitch, Lasse Hallenga, Sarya Derado, Kelly T. Hughes, Philipp F. Popp, Emmanuelle Charpentier, Yann S. Dufour, Marc Erhardt

## Abstract

Many bacteria assemble multiple flagella, although building flagella imposes a substantial biosynthetic and energetic cost. We used the peritrichously flagellated model organism *Salmonella enterica* to quantify how flagellar abundance affects bacterial growth, proteome allocation, and motility. For this, we generated genetically modified strains with inducible or constitutive expression of the flagellar master regulator *flhDC*, resulting in a panel of strains ranging from nearly non-flagellated to hyperflagellated cells. We found that higher flagellar investment reduced growth rate and redirected proteome allocation, with an expansion of the flagellar sector occurring largely at the expense of the ribosomal sector. Growth analyses of flagellar assembly mutants, combined with cost modeling, suggested that flagellin biosynthesis dominated the energetic burden, whereas motor rotation contributed a smaller additional cost. Despite the associated cost, increased flagellation improved soft-agar spreading, single-cell swimming speed, effective diffusivity, and competitive fitness in spatially structured environments. A coarse-grained proteome-allocation model parametrized from these data reproduced the observed growth penalties, while simulations of navigation in dynamic chemical gradients predicted that motility benefits saturate near a flagellar investment of 3% of proteome mass. Beyond this point, rising biosynthetic costs outweigh diminishing motility gains. In summary, these results support a quantitative cost-benefit model in which heterogeneous, spatially structured environments favor an intermediate number of flagella by balancing motility benefits against the biosynthetic costs of building and operating multiple flagella.

## Introduction

Bacteria move through their environments using diverse strategies, including flagella-mediated swimming, type IV pili-mediated twitching, and gliding motility ^1,2^. Among these, flagellar swimming is one of the most widespread and best characterized, and in many species it is a major determinant of fitness because it promotes dispersal, access to nutrients, avoidance of harmful conditions, and colonization of host niches and surfaces ^3^. The bacterial flagellum is a large extracellular rotary nanomachine that propels cells by rotating a helical filament. In peritrichous species such as *Escherichia coli* and *Salmonella enterica*, multiple flagella distributed over the cell surface act together to support swimming and chemotaxis ^4^. However, these benefits come at a substantial cost. Assembly of a single flagellum requires the coordinated expression of dozens of genes, the incorporation of tens of thousands of protein subunits, and continuous rotation consumes the ion motive force that otherwise drives metabolic processes ^4–6^. For *E. coli*, the combined cost of producing and operating flagella has been estimated at between 2% and 10% of the cell’s total biosynthetic and energetic budget ^7^. Because flagella are also exposed surface structures, they can increase susceptibility to recognition by the host immune system ^8^ and flagellotrophic bacteriophages ^9^. Flagellar motility is therefore not merely a locomotive trait but a complex cellular investment whose value depends on ecological context. This substantial cost explains why bacteria have evolved elaborate transcriptional and post-translational control of flagellar synthesis ^6,10,11^. Flagellar morphology, location and number differ markedly across bacterial species ^12^. Some species, such as *Vibrio cholerae*, carry a single polar flagellum, whereas others, such as *Bacillus subtilis*, distribute dozens of flagella over the cell body. Bacteria with a single polar flagellum are most often associated with aqueous habitats, whereas multi-flagellated species are more commonly found in more viscous environments and on surfaces ^12^. Peritrichous Enterobacteriaceae such as *E. coli* and *S. enterica* assemble on average of three to six flagella per cell, yet they are neither the fastest swimmers at low viscosity nor the most proficient swarmers on wet surfaces ^2,13,14^, and their swimming behavior has been reported to be largely insensitive to variation in flagellar number ^15,16^. It is therefore not immediately evident why these species maintain this intermediate number of flagella.

Predicting the benefit of any given flagellation pattern is complicated because flagella can serve functions beyond motility, including surface attachment, biofilm formation and pathogenesis ^3,8,17,18^. Nonetheless, building and operating flagella imposes a clear cost: in batch culture, where motility confers no advantage, non-motile mutants arise repeatedly, indicating a fitness advantage ^19,20^. Further, the synthesis of tens of thousands of flagellin monomers has imposed selective pressure to bias flagellin composition toward energy-efficient amino acids ^20^. The metabolic investment in flagellar motility thus creates a fitness trade-off that should, in principle, define an optimal level of flagellar investment for each lineage and environment ^21–24^.

A quantitative framework for this trade-off requires linking three measurable layers: biosynthetic investment (the proteome mass or synthesis flux devoted to the flagellar machinery), flagellar output (flagellar number and motor function), and the fitness consequences (growth and competition under defined ecological tasks) ^24,25^. In enteric bacteria, peritrichously flagellated Enterobacteriaceae, global proteome allocation follows quantitative growth laws, in which the ribosomal sector scales with growth rate and investment in any non-essential sector is expected to reduce the resources available for translation and growth ^26,27^. Recent work in *E. coli* has begun to couple swimming physics directly to fitness cost, predicting when cells should invest more or less in motility ^24,25^. However, the precise trade-off between flagellar number, biosynthetic burden, and performance in complex environments has not been quantitatively measured in *Salmonella*. Here, we define the cost-benefit trade-off of peritrichous flagellation in *S. enterica* serovar Typhimurium by tuning *flhDC* expression with inducible and constitutive synthetic promoters and quantifying the effects on growth, proteome allocation, motility, and competition. Combining single-cell imaging, proteomics, swimming assays, structured-environment fitness measurements, and coarse-grained modelling, we show that intermediate flagellar investment can balance motility gains against growth-limiting biosynthetic costs.

## Results

### Inducible and synthetic promoters tune flagellar assembly across a broad range

To vary flagellar assembly, we replaced the native class 1 promoter of *flhDC* in *S. enterica* serovar Typhimurium LT2 using two complementary strategies. An AnTc-inducible P*tetA*-*flhDC* allele enabled dose titration within one genetic background, whereas constitutive synthetic promoters provided fixed, inducer-free expression for prolonged assays. The inducible construct placed expression of the master regulator FlhD4C2 under AnTc control ^28,29^ (Figure 1A). All strains carried the surface-exposed FlgE-S171C substitution, which was covalently labelled with fluorophore-coupled maleimide to produce discrete hook foci ^29^. Because each hook marks a completed hook–basal body, hook counts were used as a proxy for flagellar number. Filaments were detected separately by anti-FliC immunostaining, and TH9677, carrying FlgE-S171C and the native *flhDC* promoter, served as the WT comparison (Figure 1B). Hook number increased dose-dependently and plateaued at high AnTc concentrations (Figure 1C). Mean hook counts were 0.06 ± 0.05 without AnTc and 1.4 ± 0.4 at 0.25 ng/mL. Counts rose to 2.3 ± 0.4 at 0.5 ng/mL and 3.4 ± 0.1 at 1 ng/mL, and reached their maximum at 3.7 ± 0.3 and 3.7 ± 0.5 for 2 and 4 ng/mL, respectively. For comparison, WT cells carried 2.6 ± 0.6 hooks. At 0.5 ng/mL, mean hook and filament counts approximated WT values (2.3 versus 2.6 hooks and 2.3 versus 2.3 filaments). Filament counts broadly followed the hook response, reaching 3.7 ± 0.3 at 2 ng/mL before declining to 3.3 ± 0.7 at 4 ng/mL (Figure 1D). Matched-cell analysis showed a strong association between hook and filament number (Figure 1E). Among 18,873 cells, 1,857 (9.84%) had more hooks than filaments, whereas none had more filaments than hooks. This directional mismatch is consistent with a minority of hook–basal bodies failing to switch to late-substrate secretion and consequently not assembling a filament ^29^.

**Figure 1.**
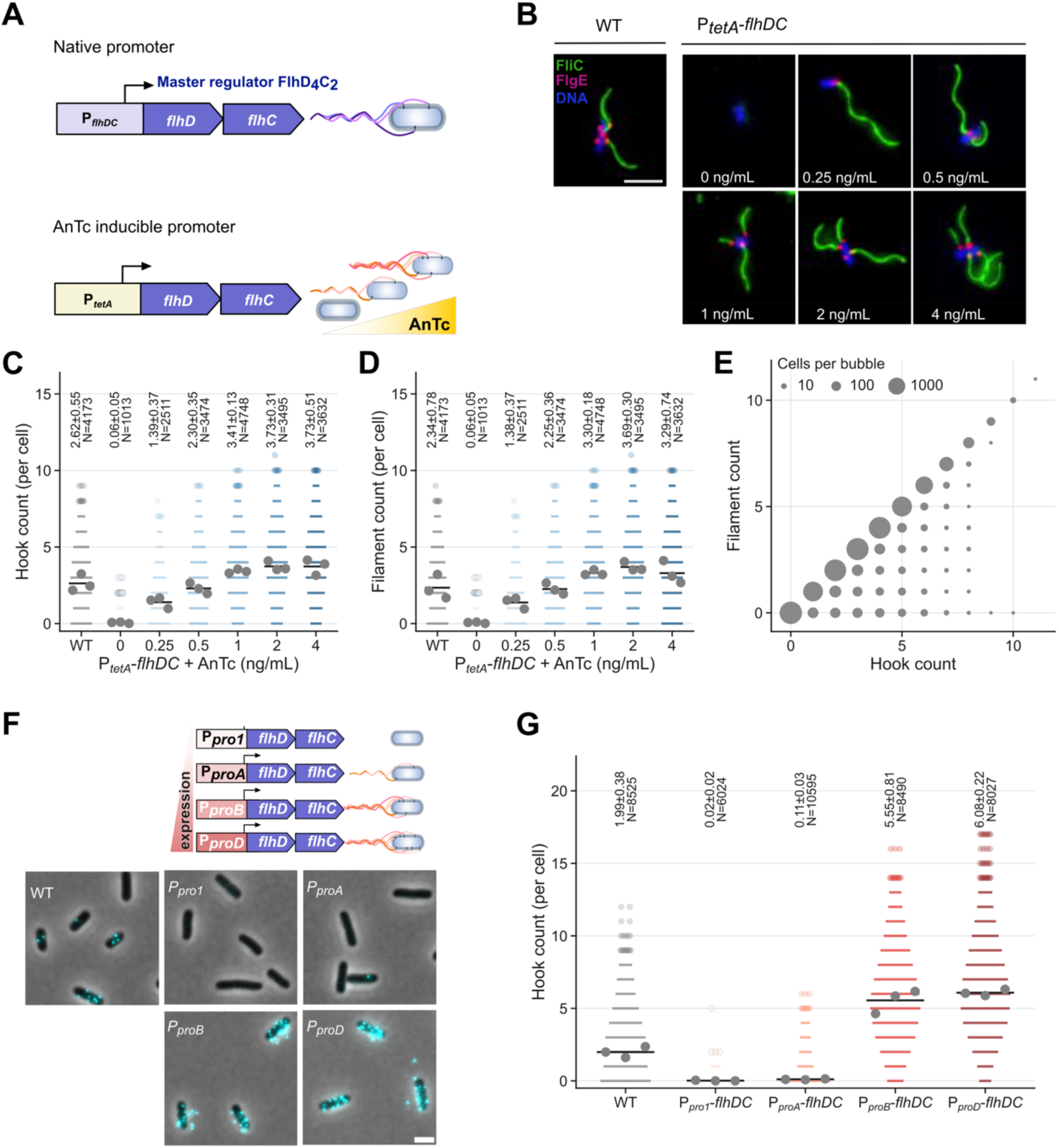
Tunable control of flagellar number per cell. A, Inducible control of flagellar number. The native class 1 promoter of *flhDC* was replaced by an anhydrotetracycline (AnTc)-inducible promoter, linking FlhD4C2 abundance and flagellar number to AnTc concentration. B, Fluorescence micrographs of cells grown with 0 - 4 ng ml^−1^ AnTc. Filament (FliC), green; hook (FlgE-S171C), magenta; DNA, blue. Scale bar, 2 µm. C, D, Hook (C) and filament (D) counts per cell from three independent experiments. For bins containing more than 12 cells, mark half-width scales with the square root of cell number; smaller bins are shown as individual cells. Grey dots are experiment means and the black bar is their mean. Labels give the mean ± s.d. across experiments and the total number of cells. E, Hook and filament counts in matched cells. Bubble area scales with the square root of cell number. F, Constitutive control of flagellar number using four synthetic *flhDC* promoters of increasing strength. Hook signal, cyan. Scale bar, 2 µm. G, Hook counts across the promoter series, displayed as in C. WT is strain TH9677, which carries FlgE-S171C for maleimide labelling. Color indicates AnTc concentration or promoter strength. C, D and G were compared with WT by two-sided Welch’s *t*-tests on the three experiment means, with Benjamini-Hochberg correction within each panel; exact *P* and *q* values are provided in Source Data.

For inducer-free control, we replaced the native *flhDC* promoter with the synthetic promoter series P*pro1*, P*proA*, P*proB* and P*proD*. These promoters share the same −35 element but carry distinct −10 elements with previously characterized strengths ^30^ (Figure 1F). Hook output increased in a step-like manner as expected (Figure 1G). WT cells in this experiment carried 2.0 ± 0.4 hooks. P*pro1*-*flhDC* and P*proA*-*flhDC* produced 0.02 ± 0.02 and 0.11 ± 0.03 hooks per cell, respectively. Correspondingly, 99.6% of P*pro1*-*flhDC* cells and 93.2% of P*proA*-*flhDC* cells lacked detectable hooks. P*proB*-*flhDC* and P*proD*-*flhDC* increased hook counts to 5.6 ± 0.8 and 6.1 ± 0.2 per cell, respectively. Together, the inducible and constitutive systems spanned near-zero to more than six completed hook–basal bodies per cell, providing complementary control of flagellar assembly.

### Growth rate decreases as flagellar production increases

To quantify the growth consequences of flagellar production, we measured batch-culture growth rates across the inducible and constitutive promoter series. Cells were grown in minimal medium supplemented with 1% (w/v) yeast extract, conditions chosen to favor flagellar gene expression by reducing RflP-dependent inhibition of FlhD4C2 ^31,32^. Cultures were transferred from exponential phase to minimize variation associated with entry into growth. Relative to the same-day WT, uninduced cultures and cultures induced with 0.25 ng/mL AnTc grew 8.3% and 6.4% faster, respectively. By contrast, cultures induced with 1, 2, or 4 ng/mL grew 6.3%, 7.2%, and 6.4% more slowly, respectively (Figure 2A). At 0.5 ng/mL AnTc, the mean growth rate did not differ detectably from the WT. Although growth rates of individual experiments were heterogeneous, potentially reflecting coexisting flagellated and non-flagellated subpopulations, the mean hook number per cell matched the WT. Thus, 0.5 ng/mL AnTc produced hook-number and growth phenotypes consistent with native *flhDC* output. The heterogeneous growth rates could reflect differences among cells in the timing of FliA-dependent class 3 gene expression, which begins only after completion of the hook– basal body and relief of FlgM-mediated inhibition ^28,33^. The constitutive promoter series produced a comparable separation between low and high flagellar output. The weak P*pro1*-*flhDC* and P*proA*-*flhDC* strains, which produced very few hooks, grew 16.1% and 17.6% faster than the WT. The strong P*proB*-*flhDC* and P*proD*-*flhDC* strains, which produced approximately 5.6 and 6.1 hooks per cell, grew 7.8% and 12.2% more slowly, respectively (Figure 2B). These results thus identify a growth cost associated with high flagellar output.

**Figure 2.**
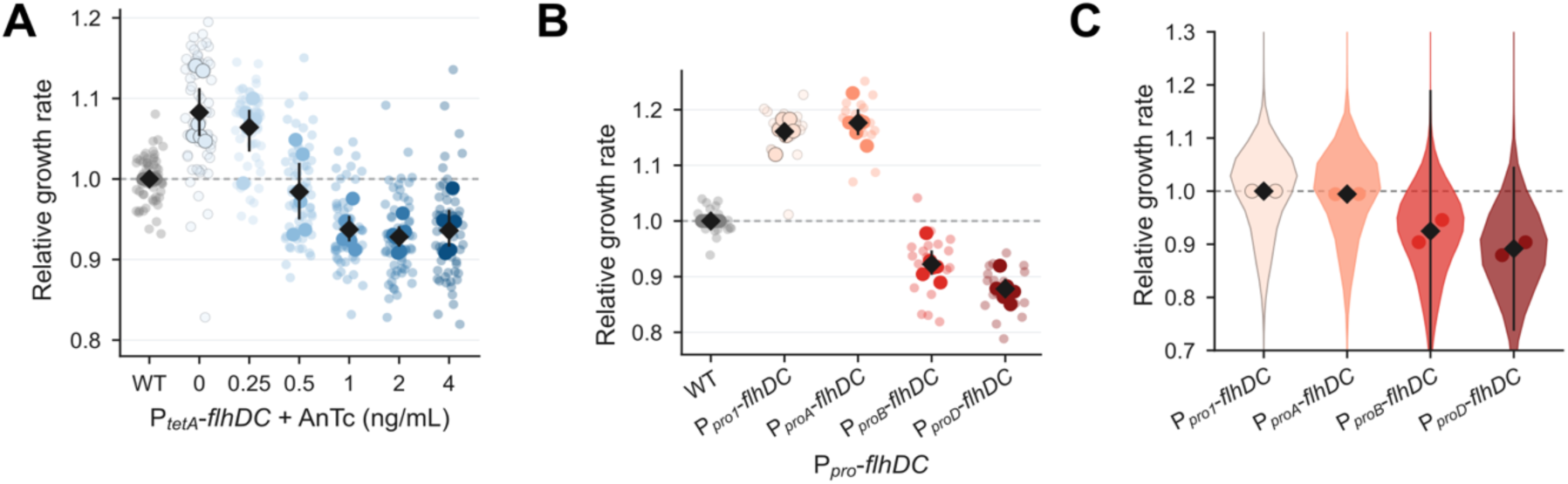
Growth rate decreases with flagellar number. A, B, Batch-culture growth rates across the AnTc (A) and promoter (B) series, normalized to the same-day WT mean. A contains ten cultures per condition on each of six days; B contains three cultures per condition on each of six days. Pale marks are individual cultures, filled marks are day means and the black diamond is the mean of the six day values with its 95% confidence interval. The experiment day is the unit of analysis. C, Single-cell growth in a microfluidic mother machine, normalized to the P*pro1*-*flhDC* mean of the same experiment. The violin shows the pooled growth rate distribution from 126,934 division cycles and the two points are the means of two independent experiments per strain. Color indicates AnTc concentration in A and promoter strength in B and C. A and B were analyzed using two-sided paired *t*-tests on the six matched day means, with Benjamini-Hochberg correction within each panel; exact *P* and *q* values are provided in Source Data.

We next asked whether the separation between weak and strong promoters was also evident at the single-cell level. Growth rates were measured in a microfluidic mother machine from successive cell-division cycles (Figure 2C, Supplementary Figure S1, Supplementary Figure S2 and Supplementary Table S1). After normalization to the replicate-matched P*pro1*-*flhDC* mean, P*proA*-*flhDC* grew approximately 0.5% slower, P*proB*-*flhDC* approximately 7.5% slower, and P*proD*-*flhDC* approximately 10.8% slower. The single-cell distributions therefore showed the same flagella-dependent growth rate phenotype as the batch-culture measurements, with slower growth in the strong-promoter strains. Together, the results motivated the proteomic and modelling analyses below, which test whether high flagellar investment competes with growth-related functions for a finite protein budget ^26,27,34^.

### Flagellin production dominates the growth cost of flagellar assembly

To determine which steps of flagellar synthesis and operation impose the observed growth cost, growth rates were measured for a series of strains in which flagellar assembly was arrested at successive stages: Δ*flhDC*, which lacks downstream flagellar gene expression; Δ*flgE,* which assembles the basal body but no hook; Δ*flgM flhA*Δc, which produces flagellin but cannot secrete it; Δ*flgKL*, which secretes flagellin but cannot assemble a filament ^35^; and *motB*(D33N), which assembles flagella with non-rotating motors ^36^ (Figure 3A).

**Figure 3.**
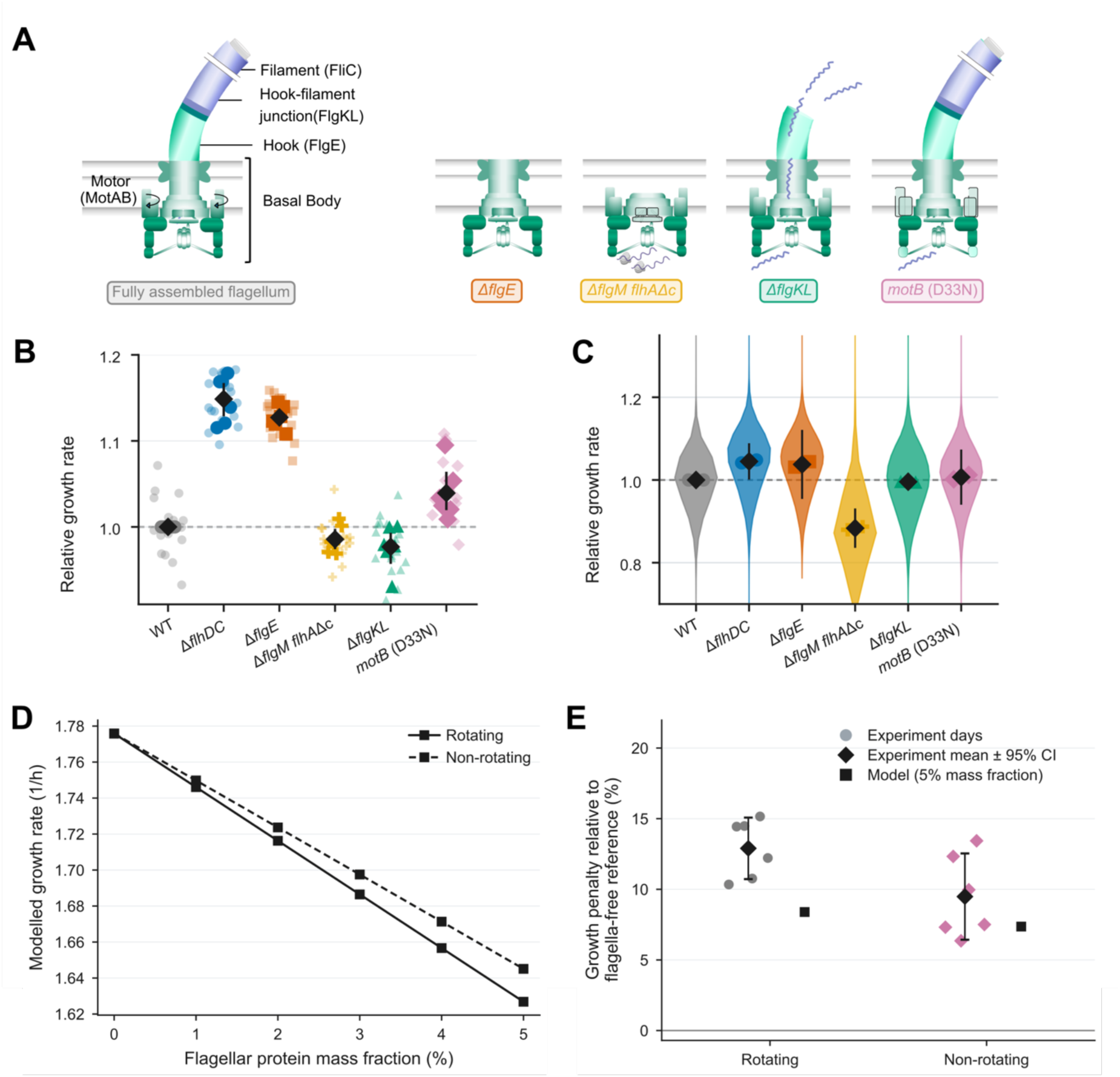
Flagellin production dominates the growth cost. A, Mutants that arrest flagellar assembly at defined stages. Δ*flhDC* lacks downstream flagellar gene expression; Δ*flgE* assembles the basal body but no hook; Δ*flgM flhA*Δc produces flagellin without secreting it; Δ*flgKL* secretes flagellin without assembling a filament; and *motB*(D33N) assembles a non-rotating flagellum. B, Batch-culture growth of the mutant series, normalized to same-day WT. Three cultures were measured per strain on each of six days; display layers are as in Figure 2A. C, Single-cell growth of the same strains measured in two independent mother-machine experiments. The violin shows the pooled growth rate distribution of 110,983 division cycles. Colors and symbols identify strains as in A and B. D, Growth rate predicted by the cell-economy model (Figure 4A) as a function of flagellar protein allocation for rotating (solid) and non-rotating (dashed) flagella. E, Measured growth penalty relative to the flagella-free Δ*flhDC* strain. Points are experiment-day values, the black diamond is their mean with a 95% confidence interval and the black square is the model prediction at 5% flagellar allocation. B and E were analyzed using two-sided paired *t*-tests on six matched day values, with Benjamini-Hochberg correction within each panel; exact *P* and *q* values are provided in Source Data.

The non-flagellated Δ*flhDC* strain grew fastest, at 14.9% above the same-day WT (Figure 3B). Δ*flgE*, which assembles only the basal body, grew almost as fast, at 12.7% above WT, indicating that early flagellar assembly contributed a comparatively small fraction of the total cost. By contrast, strains retaining flagellin protein production grew at approximately WT rates. Δ*flgM flhA*Δc, which expresses flagellin without secreting it, grew 1.4% slower than the WT, and Δ*flgKL*, which secretes flagellin without assembling a filament, grew 2.3% slower. Because these strains produce basal-body numbers comparable to the WT, flagellin expression accounts for essentially the whole growth cost, whether or not the protein is secreted or assembled, which identifies flagellin production as the dominant biosynthetic burden. The single-cell mother machine measurements provided support for this ordering (Figure 3C). Δ*flhDC* and Δ*flgE* grew faster than WT, whereas Δ*flgKL* remained approximately WT-like. However, Δ*flgM flhA*Δc grew 11.7% slower than WT at the single-cell level, despite exhibiting a WT-like growth rate in batch culture. Its single-cell growth-rate distribution was unusually broad and asymmetric: the coefficient of variation was 14.6%, compared with 6.8-9.0% for the other strains, and the difference from WT ranged from −5.7% at the 95th percentile to −16.5% at the 5th percentile. This heterogeneity could reconcile the single-cell and population-level measurements, because faster-growing cells contribute disproportionately to the expansion of a batch culture and can therefore mask a slower-growing subpopulation. The broad distribution may reflect an uneven cellular burden caused by constitutive flagellin production in the absence of an export route.

Rotation added a much smaller cost. The non-rotating *motB* (D33N) mutant grew 3.9% faster than the WT in batch culture. Measured against the flagella-free Δ*flhDC* reference, the WT carried a growth penalty of 12.9% and the non-rotating mutant one of 9.5% (Figure 3B). Abolishing rotation therefore relieved 3.4 percentage points of the growth rate costs associated with operating flagella, corresponding to approximately 26% of the total cost measured for WT flagella. To compare the measured cost of motor rotation with an energetic prediction, we used the coarse-grained cell-economy model described in more detail below (Figure 4A). The model distributes a limited protein pool among eight cellular sectors, such that increased flagellar allocation reduces the resources available to other functions ^27,37^. For the present comparison, growth was predicted at fixed flagellar mass fractions with the energetic demand of motor operation either included or omitted. The rotating condition incorporated an ATP-equivalent demand for the proton-driven motor, whereas this demand was set to zero in the non-rotating condition ^7,36,38^. Simulated growth declined progressively with increasing flagellar mass fraction under both conditions. At a flagellar mass fraction of 5%, the predicted growth penalty relative to the non-flagellated state was 8.4% with rotation and 7.4% without rotation (Figure 3D). Rotation therefore contributed 1.0 percentage point to the predicted penalty, compared with 3.4 percentage points in the batch culture experiment. Despite this quantitative difference, both approaches supported the same cost hierarchy: the principal growth penalty arose from flagellar protein production, whereas motor operation imposed a smaller additional cost. This relative weighting differs from the bioenergetic estimate of Schavemaker and Lynch, who attributed 5.2% of the whole-cell energy budget to operating *E. coli* flagella and 5.0% to their construction ^7^.

**Figure 4.**
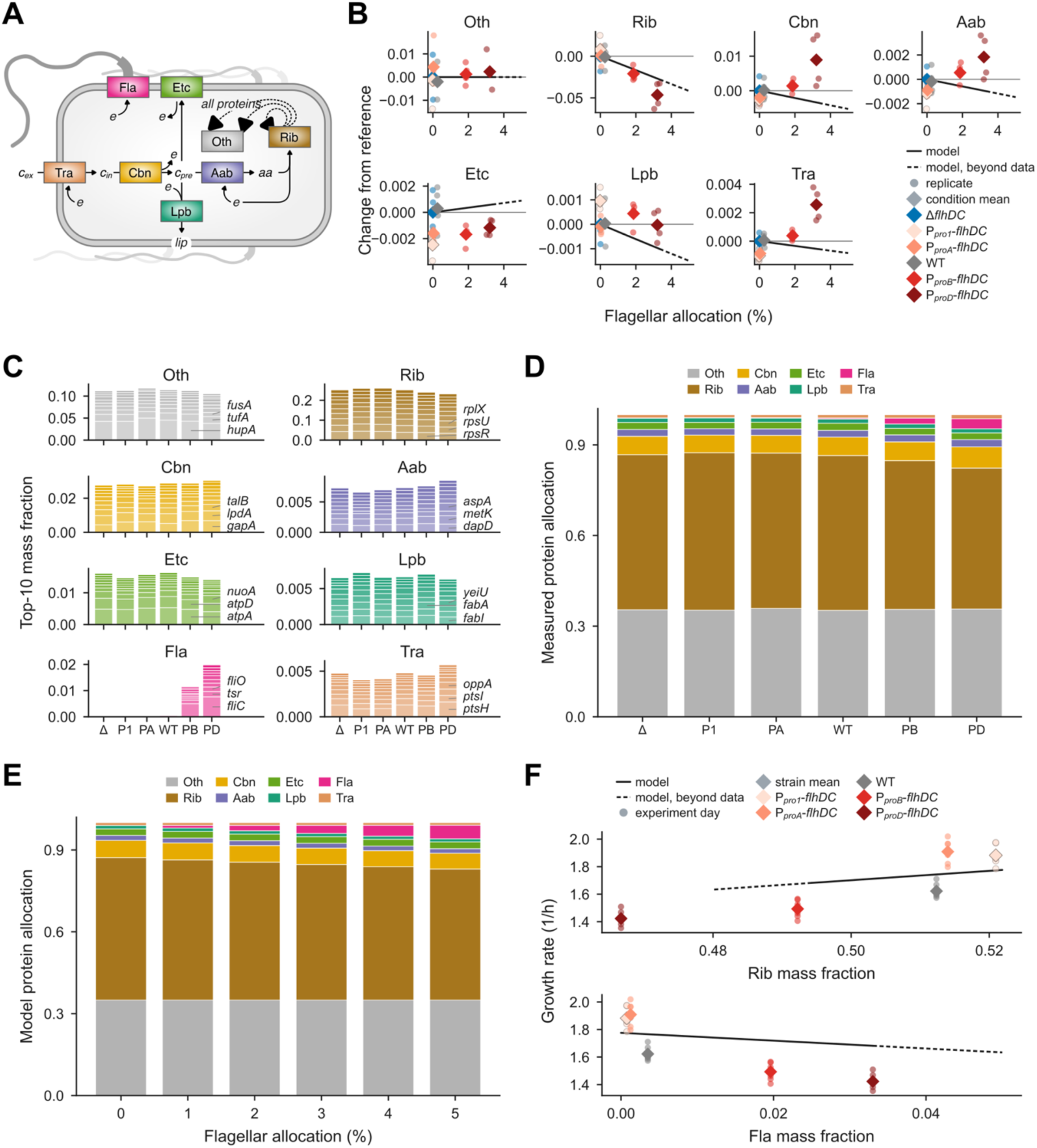
Proteome allocation under increasing flagellar investment. A, Cell-economy model in which eight sectors compete for a limited protein budget: flagella (Fla), ribosomes (Rib), transport (Tra), carbon metabolism (Cbn), amino-acid biosynthesis (Aab), lipid biosynthesis (Lpb), electron transport (Etc) and other proteins (Oth). B, Sector changes with increasing flagellar allocation, measured by mass spectrometry in four biological replicates per strain and referenced to Δ*flhDC*. Small circles are replicates and diamonds are strain means. Lines show model output, solid over the measured range and dashed above 3.34% flagellar allocation. Across the six strain means, ribosomal allocation decreased with flagellar allocation (slope, −1.49; 95% confidence interval, −1.81 to −1.17; ordinary least squares, two-sided *P* = 0.00021; *q* = 0.0015 after correction across seven non-flagellar sectors). C, Protein contributions of the top 10 proteins by mass within each sector, averaged across four biological replicates. Sector totals are given in D. D, Measured sector composition by strain. E, Modelled sector composition with increasing flagellar allocation. D and E use the same sector colors. F, Growth rate against ribosomal and flagellar allocation. Small circles are six growth-experiment days per strain and therefore share the corresponding strain-level proteomics value; diamonds are strain means. Lines show model output and were not fitted to these data. Sector assignment, protein-labelling criteria and remaining regressions are described in Methods and Source Data.

### Increasing flagellar investment reduces ribosomal allocation and growth

To determine how the biosynthetic cost of flagellation is distributed across the proteome, we performed mass-spectrometry-based proteomics on the synthetic-promoter strains P*pro1*-*flhDC*, P*proA*-*flhDC*, P*proB*-*flhDC* and P*proD*-*flhDC*, together with WT and Δ*flhDC* controls, using four biological replicates per strain. From the 4,533-entry reviewed UniProt reference proteome, we quantified 2,751 protein accessions. Of these, 1,304 protein groups could be assigned to KEGG-derived pathways and together represented 70% of the measured protein mass. We grouped the quantified proteome into eight sectors - flagella, ribosomes, transport, carbon metabolism, amino-acid biosynthesis, lipid biosynthesis, electron transport and a residual “other” sector - corresponding to the coarse-grained “super-enzymes” of the cell-economy model (Figure 4A). In this framework, the sectors compete for a limited cellular protein pool, linking flagellar investment to allocation elsewhere in the proteome ^27,37^.

The dominant response across the strain series was an inverse relationship between flagellar and ribosomal allocation (Figure 4B,D). Across the six measured conditions, ribosomal allocation decreased with flagellar allocation. The flagellar sector increased to 3.3% of total protein mass in P*proD*-*flhDC*, whereas the ribosomal sector decreased from 51.3% to 46.7%. This redistribution was not restricted to the flagellar and ribosomal sectors: carbon metabolism, amino-acid biosynthesis and transport showed smaller but detectable increases, whereas the other, electron-transport and lipid-biosynthesis sectors showed no detectable trends. Increased flagellar production therefore caused a broader, but strongly ribosome-dominated, redistribution of the proteome. At the individual-protein level, FliC showed the largest increase between Δ*flhDC* and P*proD*-*flhDC*, consistent with the tens of thousands of flagellin subunits required to construct each filament ^5,39^. Smaller increases occurred across structural, motor, regulatory and chemotaxis proteins, including FliO, FlgE, MotA, FlhD, CheA, CheW, Tsr and Tar (Figure 4C; Supplementary Figure S3). Thus, the expansion of the flagellar sector primarily reflected flagellin production but extended across the broader motility apparatus. As shown in Figure 4F, growth rate increased with ribosomal allocation and decreased with flagellar allocation. The positive ribosome-growth association is consistent with established bacterial growth laws and growth-dependent changes in the fraction of active ribosomes ^26,40^. Together with the strong inverse relationship between flagellar and ribosomal mass fractions, these data are consistent with competition for a finite proteome budget.

We next asked whether a finite-proteome constraint was sufficient to account for the measured changes. We parameterized the coarse-grained cell-economy model using the measured sector abundances together with literature-derived or explicitly estimated kinetic and structural constants (Supplementary Table S2). Increasing the imposed flagellar mass fraction produced a monotonic decline in simulated growth and a corresponding reduction in ribosomal allocation (Figures 3D and 4B,E,F). The model therefore captured the direction of the dominant flagella–ribosome trade-off. It did not, however, reproduce the complete proteome response: carbon metabolism, amino-acid biosynthesis and transport decreased in the model as it expects a lower demand for these sectors with reduced growth, while these sectors increased experimentally. The predicted increase in electron transport was not detected in the proteomics data and the experimental growth–allocation slope was approximately 4.8-fold steeper than the model prediction. The model thus recapitulates the qualitative proteome constraint imposed by flagellar investment but not the full compensatory response or the magnitude of the associated growth cost.

### Dynamic gradient simulations support an intermediate optimum of flagellar investment

Having quantified the growth cost of flagellar production, we next asked whether greater flagellar investment could be offset by improved access to nutrient-rich environments. We coupled the cell-economy model to a dynamic gradient-navigation simulation in which cells with fixed flagellar allocations of 0.5-5% moved through a glucose concentration gradient. After 8 h, cells with a 3% flagellar allocation travelled 3.7-fold further than cells with a 0.5% allocation (7.91 mm versus 2.15 mm). Raising the allocation to 5% extended this to only 4.0-fold (8.50 mm), an increase of 7.5% over the 3% value (Figure 5A). Most of the gain came early: a 1% allocation already carried cells 2.2-fold further than 0.5%, and a 2% allocation 3.3-fold further. Flagellar investment therefore produced steep initial gains in substrate access followed by diminishing returns.

**Figure 5.**
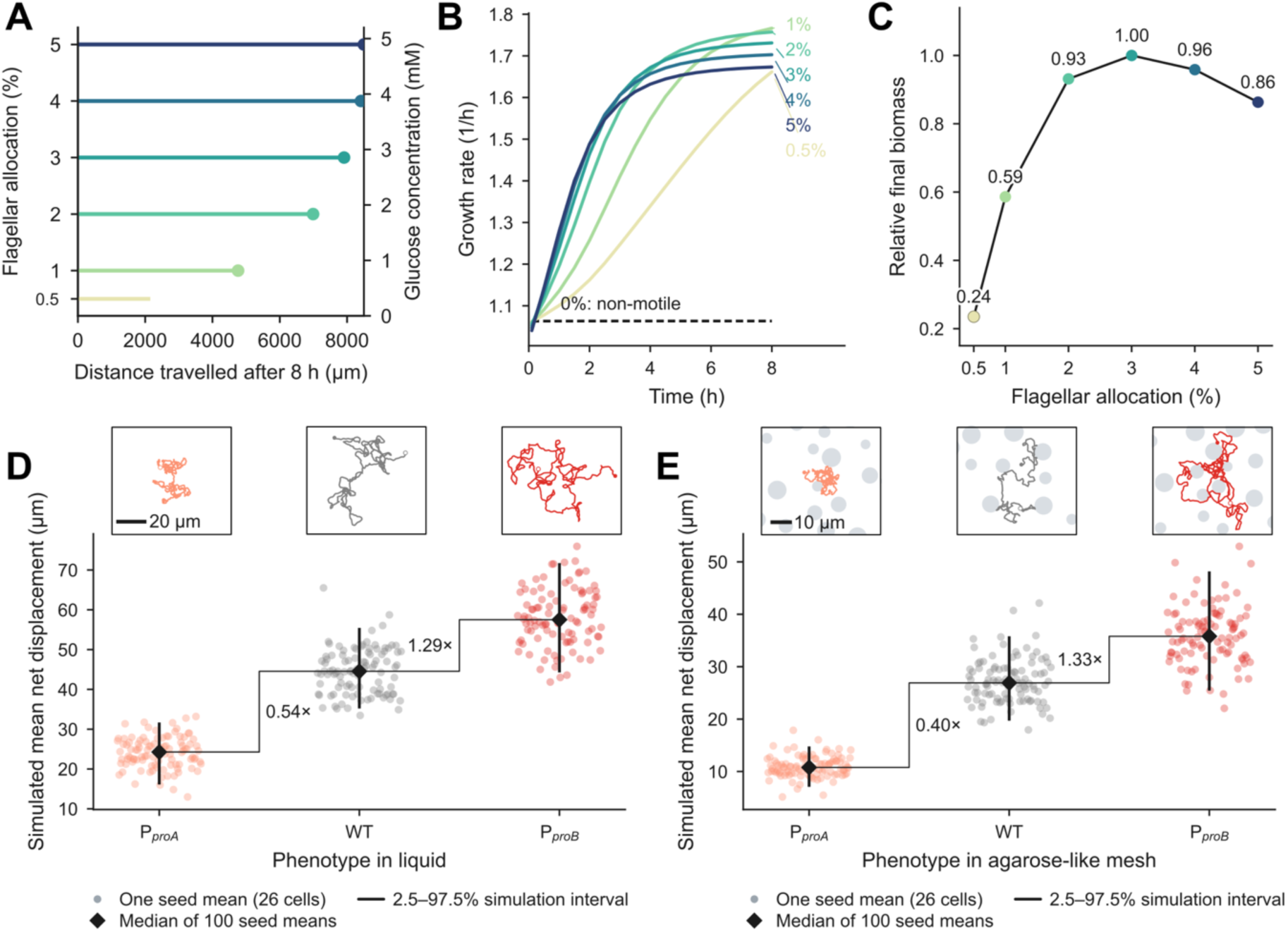
Simulated motility benefits support an intermediate optimum of flagellar investment. A, Distance travelled after 8 h by simulated cells with fixed flagellar allocations of 0.5-5%. Horizontal segments denote the distance travelled and circles mark the corresponding endpoints. Grey shading shows the spatial glucose profile established over 3 h and subsequently held fixed during the navigation simulation. B, Modelled growth-rate trajectories over 8 h. Colored lines represent flagellar allocations of 0.5–5%; the dashed grey line represents a non-motile cell with 0% flagellar allocation. C, Final biomass after 8 h, normalized to the maximum among the six evaluated allocations. D, E, Active-particle simulations of mean net displacement in liquid (D) and an agarose-like obstacle mesh (E). Each small point is the mean displacement of 26 simulated cells for one of 100 fixed random seeds. Diamonds denote the median of the 100 seed means, and intervals span their 2.5th-97.5th percentiles. The inset displays the trajectory of one representative, simulated cell. Run speed, motile fraction and persistence time were calibrated from measured motility parameters. All panels show deterministic or seeded model outputs and carry no statistical tests. Calibration and validation are described in Methods.

Improved substrate access translated into higher growth rates (Figure 5B). The non-motile reference remained 8.5 mm from the source and maintained a growth rate of approximately 1.063 h^−1^ at its initial glucose concentration of 0.091 mM. By contrast, the motile allocations reached 1.66–1.77 h^−1^ after 8 h, corresponding to increases of 56–66% over the non-motile reference. When compounded over the simulation, these differences produced final biomasses 8.3- to 35.5-fold greater than that of the non-motile cell. Among the six motile allocations evaluated, final biomass increased sharply from 0.24 of the sampled maximum at 0.5% allocation to 0.59 at 1% and 0.93 at 2%, reached its highest sampled value at 3%, and then declined to 0.96 and 0.86 of the maximum at 4% and 5%, respectively (Figure 5C). Thus, the simulations predict that fitness is maximized at an intermediate flagellar investment of approximately 3% of proteome mass, where cells capture most of the motility benefit while avoiding the disproportionate biosynthetic cost of further flagellar expression.

We next used an active-particle model parameterized with the measured run speed, motile fraction and persistence time to illustrate how the three motility phenotypes translate into dispersal in liquid and structured environments (Figure 5D,E; Supplementary Figure S4). Across 100 simulation seeds, median seed-mean net displacement in liquid increased from 24.3 µm for P*proA*-*flhDC* to 44.6 µm for WT and 57.5 µm for P*proB*-*flhDC*. Introducing an agarose-like obstacle mesh reduced displacement in all three phenotypes, to 10.8, 26.9 and 35.8 µm, respectively, with the largest proportional reduction occurring in P*proA*-*flhDC*. Representative trajectories likewise showed greater confinement and trapping of P*proA*-*flhDC* cells, whereas WT and particularly P*proB*-*flhDC* cells explored a larger area (Supplementary Figure S4). Thus, the model translates the measured motility differences into a greater separation of dispersal capacity in a porous environment, consistent with evidence that physical confinement reorganizes bacterial dispersal ^41^.

### Making multiple flagella is beneficial for motility in complex environments

To determine whether the motility benefit of increased flagellar production offsets its growth cost in a structured environment, we measured radial expansion through 0.3% motility agar. In the inducible P*tetA*-*flhDC* series, mean halo diameter increased progressively with AnTc concentration. Relative to WT, expansion rose from 3% without AnTc to 46%, 82% and 99.6% at 0.25, 0.5 and 1 ng ml^−1^ AnTc, respectively. It then plateaued at 111% at 2 and 4 ng ml^−1^ AnTc (Figure 6A). This plateau coincided with the saturation of hook and filament numbers observed over the same induction range (Figure 1). We next examined the constitutive promoter series in a Δ*rflP* background, which removes nutrient-dependent post-translational regulation of FlhDC ^31,42^. Relative halo diameter was 4% of WT for P*pro1*-*flhDC* and 79% for P*proA*-*flhDC*. Notably, the P*proD*-*flhDC* strain, despite carrying the most flagella, was slightly less motile than the P*proB*-*flhDC* strain. Expansion increased to 174% for P*proB*-*flhDC* but declined to 161% for P*proD*-*flhDC* (Figure 6B). Thus, maximal flagellar production did not maximize halo expansion. The lower expansion of P*proD*-*flhDC* relative to P*proB*-*flhDC*, despite its greater hook number, is consistent with the increased growth burden of flagellar overproduction observed in Figure 2.

**Figure 6.**
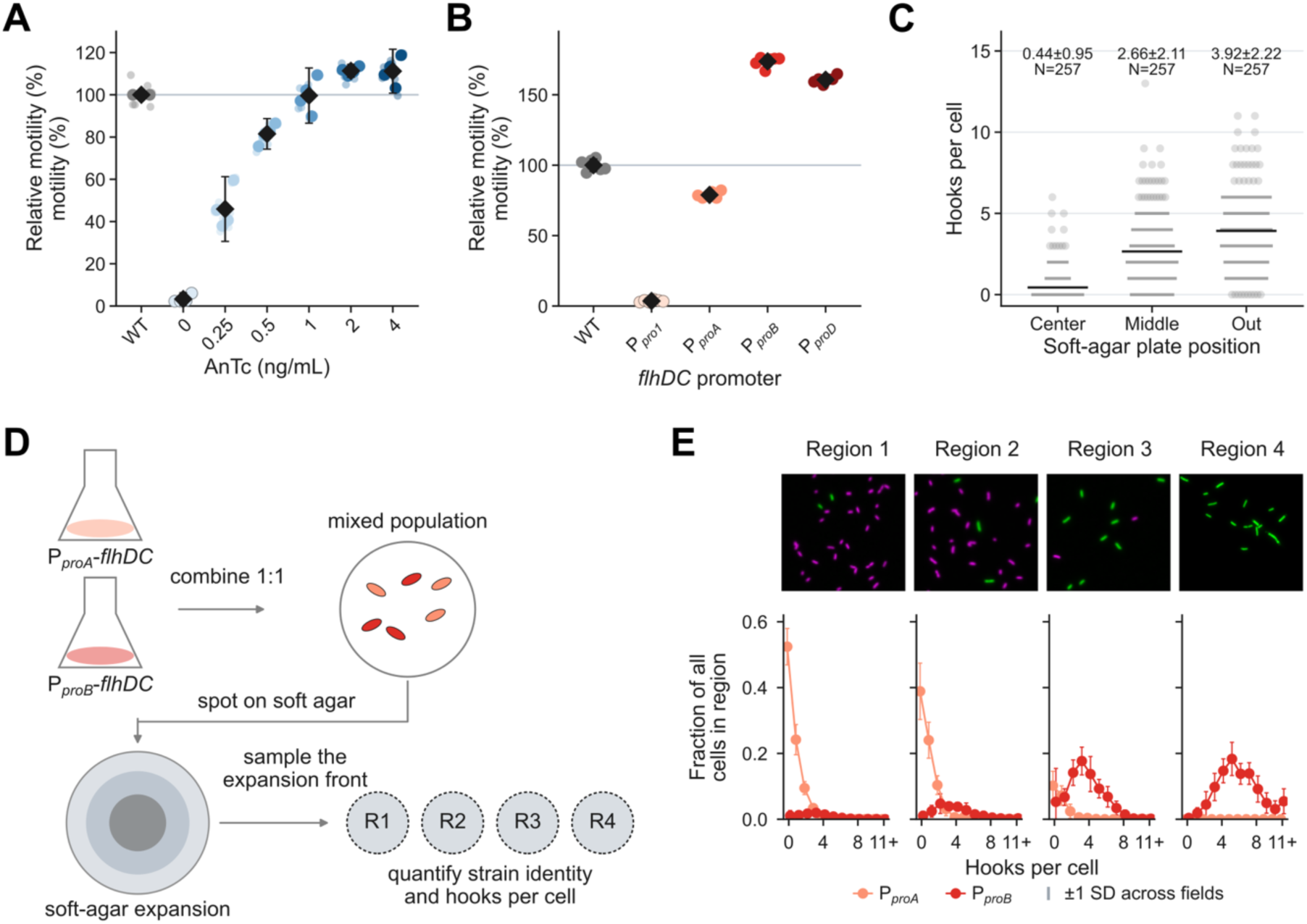
Flagellar production promotes soft-agar expansion and spatial sorting in a structured environment. A, B, Soft-agar motility of the P*tet-flhDC (A) and Ppro*-*flhDC* (B) series, measured as halo diameter in 0.3% agar and normalized to WT. A, AnTc series, comprising 93 independent motility halo measurements across four experiment days. B, Promoter series, comprising one independent halo measurement per strain on each of six days. Pale marks are individual halo measurements, filled marks are experiment-day means and the black diamond is the mean of the experiment-day values with its 95% confidence interval. Strains in B were compared with WT using two-sided paired *t*-tests on six matched halo measurements; exact *P* values are provided in Source Data. C, Hook counts at the center, middle and edge of one soft-agar halo, with 257 cells per position, displayed as in Figure 1C. D, Competition assay. P*proA*-*flhDC*-mScarlet and P*proB*-*flhDC*-eGFP cells were mixed 1:1, inoculated into soft agar and recovered from regions R1 - R4 between the inoculation point and halo edge. E, Representative fluorescence images and hook-count composition across R1 - R4, comprising 8,581 cells. Points are mean field-level fractions of cells with the indicated hook count; error bars are ±1 s.d. across 18, 13, 13 and 12 imaging fields for R1 to R4, respectively, from one representative experiment.

To test whether cells with more flagella are spatially enriched also under native regulatory control, we quantified hook number as a proxy for flagellar number at defined radial positions within a motility halo of the WT strain. Mean hook number increased from 0.4 ± 1.0 at the center to 2.7 ± 2.1 in the middle and 3.9 ± 2.2 at the outer position (Figure 6C). Correspondingly, the fraction of cells without hooks decreased from 76% at the center to 14% in the middle and 4% at the outer position. Finally, we tested whether strains with different flagellar phenotypes became spatially sorted during direct competition. Equal-density cultures of P*proA*-*flhDC*-mScarlet and P*proB*-*flhDC*-eGFP were mixed, inoculated into soft agar and sampled from four regions between the inoculation site and expansion front (Figure 6D). Among 8,581 measured cells, the P*proA*-*flhDC* fraction decreased from 91% in R1 to 78% in R2 and 21% in R3, before becoming undetectable in R4. Conversely, all 870 cells recovered from R4 were P*proB*-*flhDC* and carried an average of 5.8 hooks per cell (Figure 6E). Within this plate, the more highly flagellated P*proB*-*flhDC* strain therefore occupied the expansion front despite its slower growth. This spatial sorting is consistent with a motility advantage outweighing the growth cost during soft-agar expansion.

### Higher flagellar abundance is associated with faster swimming and greater effective diffusivity in liquid and structured media

To determine how flagellar abundance relates to single-cell motility without an imposed chemical gradient, we co-imaged pairwise mixtures of WT, P*proA*-*flhDC* and P*proB*-*flhDC* cells in an 8-µm-deep microfluidic chamber. All strains carried Δ*rflP* to prevent post-translational repression of FlhDC. Each pair was mixed at a 1:1 ratio, labelled reciprocally with mScarlet and eGFP, and imaged simultaneously in liquid or 0.3% low-melting-point agarose (Figure 7). From the resulting trajectories, we quantified swimming speed and effective diffusivity, D*eff* ^43^. We also determined the fraction of tracked cells classified as swimming and calculated the persistence-equivalent timescale τ = 2D*eff*/v^2^.

**Figure 7.**
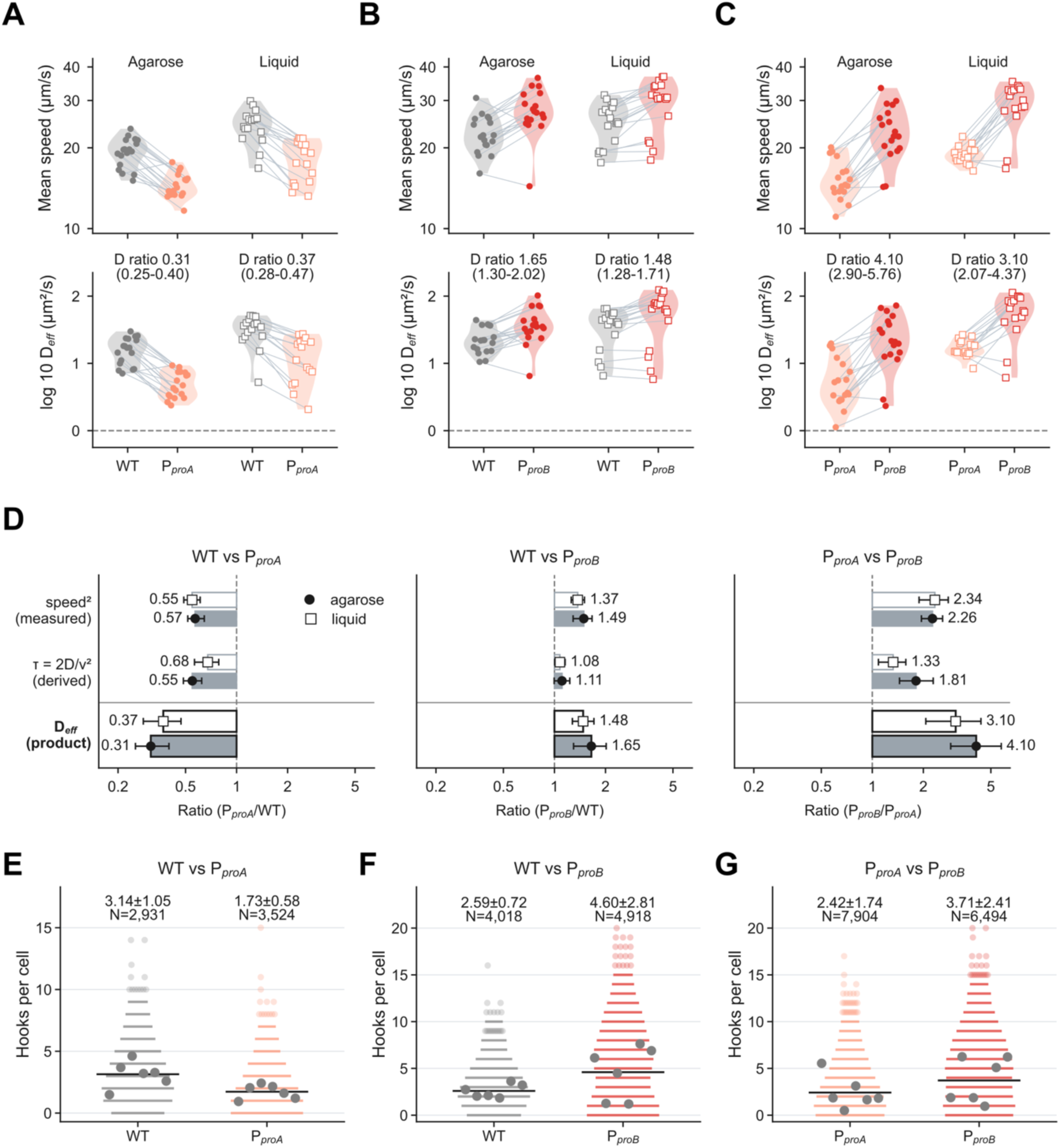
Single-cell swimming behaviour and hook number. A-C, Paired swimming measurements of WT and P*proA*-*flhDC* (A), WT and P*proB*-*flhDC* (B), and P*proA*-*flhDC* and P*proB*-*flhDC* (C). Differently labelled strains were co-imaged in an 8-µm-deep microfluidic chamber containing liquid or 0.3% low-melting agarose, with fluorophores exchanged between replicate experiments. A comprises 18 movies in agarose and 16 in liquid from six independent days; B comprises 18 movies in each medium from eight days; C comprises 18 movies in agarose and 16 in liquid from eight days. Each panel shows mean swimming speed above and mean log10 effective diffusivity (*Deff*) below. Symbols represent movies and lines connect co-imaged strains. Filled circles, agarose; open squares, liquid. Violins show the distribution across movies; headers give the medium, movie number and *Deff* ratio with its 95% confidence interval. D, Decomposition of *Deff* = *v*^2^*τ*/2 into speed squared, persistence time *τ* = 2*Deff*/*v*^2^ and their product. Estimates are equal-weight mean log ratios across paired movies; intervals are 95% paired-movie bootstrap confidence intervals. E-G, Hook counts for the corresponding strain pairs, displayed as in A-C. Grey points are six independent replicate means per strain and the black bar is their mean. Bootstrap implementation is described in Methods.

Swimming speed and effective diffusivity followed the same rank order in both media and under both reciprocal labelling configurations: P*proA*-*flhDC* < WT < P*proB*-*flhDC* (Figure 7A–C). Relative to WT, D*eff* of P*proA*-*flhDC* was 0.31-fold in agarose and 0.37-fold in liquid. Conversely, D*eff* of P*proB*-*flhDC* was 1.65-fold that of WT in agarose and 1.48-fold in liquid. The largest separation occurred in the direct comparison between the promoter strains: P*proB*-*flhDC* had 4.10-fold greater D*eff* than P*proA*-*flhDC* in agarose and 3.10-fold greater D*eff* in liquid. Paired median-speed summaries showed the same ordering, with P*proB*-*flhDC* swimming 1.64-fold faster than P*proA*-*flhDC* in agarose and 1.53-fold faster in liquid (Supplementary Figures S5 and S6).

The fraction of cells classified as swimming also differed most strongly between the low- and high-output promoters. Relative to WT, the swimming fraction of P*proA*-*flhDC* was lower by 0.27 in agarose and 0.20 in liquid. P*proB*-*flhDC* was essentially the same as WT in both media. Directly comparing the promoter strains, the swimming fraction of P*proB*-*flhDC* exceeded that of P*proA*-*flhDC* by 0.28 in agarose and 0.19 in liquid (Supplementary Figure S5). Thus, the distinction between low and high flagellar output was particularly pronounced in structured media where cells are more prone to trapping ^41^.

We note that these motility measures are related rather than independent. Effective diffusivity is calculated from cell movement, swimming cells are classified using speed and diffusivity, and τ is derived as 2D*eff*/v^2^. Both speed and τ contributed to the observed differences in D*eff* (Figure 7D), although speed explained most of the increase in P*proB*-*flhDC* relative to WT. The smaller contribution of τ was not clearly resolved in agarose. Together, these measurements describe different aspects of the same change in swimming behavior.

Hook counts from the corresponding reporter strains showed the same pairwise ordering. P*proA*-*flhDC* carried fewer hooks than WT (1.7 versus 3.1 per cell), P*proB*-*flhDC* carried more than WT (4.6 versus 2.6), and P*proB*-*flhDC* carried more than P*proA*-*flhDC* in their direct comparison (3.7 versus 2.4; Figure 7E-G). Greater flagellar abundance was therefore associated with faster swimming, a larger swimming fraction and greater effective diffusivity in both liquid and structured agarose environments.

## Discussion

Peritrichously flagellated bacteria such as *Salmonella enterica* and *Escherichia coli* commonly assemble several flagella per cell, although the number varies with strain, growth conditions and cell-cycle state ^12,24^. This variability raises a fundamental allocation problem: additional flagella may improve motility, but their synthesis and operation consume resources that could otherwise support growth. Here, we varied expression of the flagellar master regulator FlhDC over a broad range and quantified the resulting effects on flagellar assembly, growth, proteome composition and motility in *Salmonella*. The promoter replacements altered the class 1 input to the flagellar transcriptional hierarchy while retaining downstream regulatory and assembly checkpoints. Together, the results show that flagellar abundance is a tunable cellular investment whose growth cost increases with flagellar production, whereas its motility benefit depends on the physical environment and eventually saturates.

Both inducible and constitutive control of *flhDC* generated graded changes in hook and filament number, extending from predominantly non-flagellated to hyperflagellated populations (Figure 1). The strong association between hook and filament counts supports the use of labelled hooks as a proxy for flagellar number, while a minority of filament-free hooks is consistent with incomplete late-substrate switching during assembly ^29^. (Figure 1E). Across the resulting investment range, growth generally declined as flagellar output increased at both the population and single-cell levels (Figure 2). Local deviations from this trend - notably the heterogeneous phenotype at 0.5 ng/mL AnTc - indicate that the relationship is not strictly monotonic at every induction step and may be influenced by regulatory thresholds and cell-to-cell variation in assembly timing. However, the finding that the metabolic burden scales continuously with the amount of flagellar material produced is consistent with the general principle that non-essential proteome investment imposes proportional protein-budget costs ^26,44^.

Assembly-truncation mutants localized the main source of the growth burden to flagellin production (Figure 3). Removing FlhDC increased growth by 15% relative to WT, whereas a strain that assembled the basal body but no hook grew 13% faster. Basal-body assembly therefore accounted for at most approximately 2 percentage points of the total penalty under these conditions. By contrast, strains that activated late flagellar expression and produced flagellin, whether or not it was secreted or assembled into filaments, grew near or below the WT rate. Together with the proteomic observation that flagellin was the largest component of the expanding flagellar sector, these results identify flagellin-dominated class 3 expression as the principal biosynthetic burden. Motor operation imposed a smaller but measurable additional cost. Relative to the flagella-free Δ*flhDC* strain, the growth penalty was ∼13% for WT cells with rotating flagella and ∼10% for the non-rotating *motB*(D33N) mutant. Abolishing rotation therefore relieved about 3% of the measured growth penalty. The cell-economy model also predicted a small rotational contribution of 1.0 percentage point at 5% flagellar allocation. Thus, both experiment and model place biosynthesis above rotation in the cost hierarchy. This weighting notably differs from the whole-cell energy accounting of Schavemaker and Lynch, which estimated comparable construction and operating costs for *E. coli* flagella ^7^. Differences in organism, growth conditions, reference states and the quantities included in “cost” may contribute to this discrepancy.

Proteome profiling revealed how increased flagellar production was accommodated by the cell (Figure 4). In P*proD*-f*lhDC*, flagellar proteins reached ∼3% of measured protein mass, while ribosomal allocation declined from approximately 51% in the flagella-free reference to 47%. This was the dominant proteome response, but it was not the only one: carbon metabolism, amino-acid biosynthesis and transport showed smaller systematic increases, whereas the other, electron-transport and lipid-biosynthesis sectors showed no detectable trends. Flagellar investment therefore elicited broader proteome remodeling, and the inverse flagella– ribosome relationship, together with the positive association between ribosomal allocation and growth, is consistent with competition for a finite protein budget ^26,27,37^. A coarse-grained allocation model qualitatively captured the proteome-allocation trade-off. Increasing the imposed flagellar fraction reduced ribosomal allocation and growth, as observed experimentally.

Combining the modelled biosynthetic cost with the simulated benefit of gradient navigation yielded the greatest final biomass at 3% flagellar allocation among allocations sampled from 0.5% to 5% (Figure 5). This result illustrates how an intermediate optimum can emerge when the benefits of motility saturate while biosynthetic costs continue to increase. The sampled optimum is of the same order as the flagellar investment we estimate for the WT. Although the measured flagellar sector accounts for only 0.35% of protein mass in the WT, whole-cell proteomics primarily captures the intracellular flagellar pool and therefore underestimates total flagellar investment. In particular, it largely misses extracellular flagellin, both that polymerized into the external filament, which is predominantly lost during sample preparation, and exported flagellin that is not retained with the cell-associated proteome. Together, these extracellular pools constitute most of the protein mass invested in flagella. Accounting for the external filament and exported flagellin places the native investment in flagella at approximately 2-3% of protein mass for the three to four filaments per cell typical of *Salmonella* ^2,13,14^ (Supplementary Table S2). Native investment therefore approaches the modelled optimum, suggesting that native regulation may capture most of the benefits of motility while limiting the costs of overinvestment. The precise position of this optimum is nevertheless context dependent, as it varies with the imposed substrate gradient, simulation duration, the assumed relationship between motility and proteome allocation, and the parameterization of growth costs. Such context dependence is consistent with observations in *E. coli*, where nutrient conditions alter the optimal level of flagellar expression and the benefits of increased swimming speed show diminishing returns due to hydrodynamic constraints ^24^.

The experimental motility measurements support this environmental dependency (Figure 6, Figure 7). In soft agar, motility increased across most of the flagellar-number range, while the slight reduction from P*proB*-*flhDC* to P*proD*-f*lhDC* is consistent with diminishing motility gains or an increasing growth penalty at the highest investment. Hook counts increased towards the edge of a WT motility halo, and P*proB*-*flhDC* cells became enriched relative to P*proA*-*flhDC* cells during radial expansion, consistent with established growth-migration trade-offs in expanding bacterial populations ^45^ (Figure 6). Paired single-cell tracking likewise placed P*proA*-*flhDC*, WT and P*proB*-*flhDC* in increasing order of swimming performance in both liquid and agarose-containing microfluidic chambers (Figure 7). These observations were obtained without an imposed chemical gradient and therefore reflect differences in baseline trajectory behavior. These data establish a coherent association between flagellar abundance and motility phenotype, with possible contributions from altered motor switching, chemotaxis proteins or other FlhDC-dependent functions.

Active-particle simulations parameterized with experimentally measured speed, swimming fraction, and persistence predicted reduced dispersal of weakly motile cells through obstacle-rich environments (Figure 5 and Supplementary Figure S4). The predicted influence of obstacles is consistent with experimental work showing that pore confinement converts run-and-tumble motion into alternating phases of hopping and trapping ^41^. Polarly flagellated *Shewanella* can escape some traps by wrapping its filament around the cell body, illustrating how flagellar architecture may shape dispersal in structured environments ^46^. However, whether a higher number of peritrichous flagella confers a comparable advantage in intestinal mucus or other host-associated habitats remains to be tested directly in future studies.

Together, our findings establish flagellar abundance as a resource-allocation trait that couples cellular growth to motility. Greater flagellar production occurred primarily at the expense of ribosomal allocation, with the late, flagellin-dominated phase imposing most of the biosynthetic burden. Across the examined range, additional flagella enhanced motility and spatial spreading, but these gains could be offset by the associated growth cost. The fitness-maximizing allocation is therefore expected to shift with nutrient availability, gradient geometry, confinement, and the timescale over which growth and migration contribute to fitness. Our results do not define a single optimal flagellar number for *S. enterica*. Instead, they position flagellar abundance as a tunable cellular budget whose favored level emerges from an environment-dependent balance between proteome cost and motility benefit.

## Materials and Methods

### Bacterial strains and growth conditions

All strains were derived from *Salmonella enterica* serovar Typhimurium LT2 and are listed in Supplementary Table S3. Flagellar number was tuned through the master regulator *flhDC*: an anhydrotetracycline (AnTc)-inducible P*tetA*-*flhDC* allele (strain EM8242, parental TH9677) and a constitutive synthetic-promoter series of increasing strength (P*pro1* < P*proA* < P*proB* < P*proD*) replacing the native Class-I *flhDC* promoter. A *hin* fixed-orientation (FliC-ON) background, a Δ*rflP*::FRT deletion in the promoter-series and competition strains, and the *flgE*(S171C) cysteine substitution for maleimide hook labelling were common to the labelling strains. Assembly-truncation and non-rotating controls (Δ*flhDC*, Δ*flgE*, Δ*flgM flhA*Δc, Δ*flgKL*, *motB*(D33N)) and fluorescently marked competition strains (chromosomal P*rpsM*-eGFP or P*rpsM*-mScarlet-I at *amyA*) were constructed by standard methods. Bacteria were grown in Edex minimal medium (1× E salts, 0.2% D-glucose) supplemented with 1% yeast extract, shaking at 180 rpm at 37 °C or 30 °C. Batch growth curves were recorded at 30 °C; mother machine experiments were performed at 30 °C; hook and filament labelling at 30 °C; competition assay experiments at 37 °C; single-cell swimming experiments at 37 °C and soft-agar at 30 °C or 37 °C. Overnight cultures were inoculated from single colonies and diluted 1:100 for day cultures. For P*tetA*-*flhDC* induction, AnTc was added at 0.25, 0.5, 1, 2 or 4 ng/mL. Growth was monitored at OD600. Strains used in each figure are listed in Supplementary Table S4.

### Strain construction and P22 transduction

DNA fragments and selection cassettes were transferred between *S.* Typhimurium strains by generalized transduction with bacteriophage P22 HT. Phage lysates were prepared from overnight cultures in P22 broth, clarified by centrifugation and stored over chloroform. For transduction, recipient overnight cultures were mixed 1:1 with a 10^−1^ dilution of donor lysate, incubated 1 h at 37 °C and plated on selective medium; recipient-only and lysate-only mixtures served as negative controls. Transductants were purified on green-indicator plates, and phage-free colonies were identified by cross-streaking against P22 H5.

### Soft-agar motility assays

Swimming motility was assessed by inoculating 2 µL of overnight culture into soft (0.3% Bacto-agar) motility plates ^47^, incubated 3-4 h at 30 °C or 37 °C. Plates were scanned (Epson Perfection V800) and halo diameters measured in Fiji ^48^, normalized to the WT or parental strain on the same day.

### Population growth measurements and growth-rate analysis

Population growth was measured in a BioTek Synergy H1 plate reader. Day cultures grown to OD600 ≈ 0.2 were diluted to OD600 = 0.05 and 100 µL dispensed into 96-well plates; OD600 was recorded every 5 min for 16 h at 30 °C with double-orbital shaking. Each curve was processed independently: growth onset was defined as the first persistent crossing of an OD threshold (the larger of OD600 = 0.01 or the baseline median plus three times a robust estimate of the baseline noise from the first six measurements, with at least two consecutive points above threshold), and the exponential phase was identified by scanning 60-120 min windows (≥ 8 points, OD600 < 0.35) and fitting a log-linear model ln(OD600) = a + kt, evaluated by R^2^, log-scale RMSE and residual autocorrelation. The growth constant k gave the doubling time ln(2)/k. For the synthetic-promoter series k was taken from the selected window; for the assembly-mutant and P*tetA*-*flhDC* series the window was refit by nonlinear least squares to OD(t) = OD0·exp(kt). Technical replicates were averaged within experiment days, and strains or induction conditions were compared with their matched same-day reference by paired t-tests on condition-minus-reference doubling-time differences, with Benjamini–Hochberg false-discovery-rate adjustment.

### Single-cell growth tracking in a microfluidic mother machine

Single-cell growth was tracked in PDMS mother-machine chips replicated by soft lithography from a 6-inch master mold written by electron-beam lithography (Helmholtz Nano Facility, Forschungszentrum Jülich) ^49^. Following the design of Lamprecht et al. ^50^, each chip carried four cultivation channels with one inlet and two outlets, and 57 blocks of 21 single-cell traps per channel. Traps measured 30 × 1 × 1 µm with a 0.3 µm constriction at the bottom, which retained the mother cell and allowed medium to perfuse from top to bottom. Chips were cast from SYLGARD 184 (7:1 base:curing agent), cured overnight at 80 °C and bonded to high-precision cover glass by oxygen plasma (60 s, 0.8 mbar). Cells grown in Edex + 1% yeast extract at 30 °C were diluted 1:100, grown to late exponential phase and loaded into the traps. Chips were perfused with the same medium containing 0.5 mg/mL BSA under programmable pressure (MFCS, FLUIGENT). Phase-contrast images were acquired on a Nikon Eclipse Ti2 (60× Ph3 oil objective, N.A. 1.40; Hamamatsu Orca Fusion BT; Perfect Focus) at 30 °C with 90 ms exposure every 2 min for 20 h. Stacks of individually imaged mother-machine blocks were rotated and aligned with an in-house pipeline built on basic ImageJ functions ^51^. Individual traps were cropped out and their frames arranged from left to right to give kymographs, which were segmented in ilastik v1.4.0 ^52^. The resulting binary masks were processed in MATLAB v.R2022b, where each cell was detected with the regionprops function and cell length taken as the major axis length. Cells were linked into division cycles by a decision-making algorithm in MATLAB. A cycle corresponds to a cell from birth until division, death or escape from the trap. The algorithm linked the objects at the lowest kymograph position frame by frame and registered a division when the object length at t+1 fell below 90% of the length at t, terminating the cycle and initiating a new one for the bottom daughter cell. After the bottom cycles were complete, the corresponding objects were deleted from the mask and assignment continued with the objects that then occupied the lowest positions. Growth rate was estimated from the slope of a linear fit to the logarithm of cell length over the whole cycle. Estimates were retained when finite and positive, with r^2^ ≥ 0.90 and at least six length measurements per cycle. All cycles, mother-cell cycles and non-mother-cell cycles were analyzed separately. Each cycle rate was normalized to the mean growth rate of the P*pro1*-*flhDC* reference in the same experiment, calculated in sliding time windows; because the reference growth rate was stationary across the analysis window, this is equivalent to normalization by a single reference mean per experiment. Normalized rates were pooled over the analysis window (200 - 800 min after the start of imaging, 180 - 480 min in Supplementary Figure S1B) with the reference centered at 1. Growth-rate heterogeneity was quantified as the coefficient of variation of cycle-level growth rates within sliding time windows. Absolute growth rates per strain and experiment, before this normalization, are reported in Supplementary Figure S2 and Supplementary Table S1.

### Hook and filament labelling and quantification

Hooks were labelled via the S171C surface-exposed cysteine replacement mutation in *flgE* with STAR GREEN or STAR RED coupled maleimide (Abberior, 10 µM in DMF for 10 min, 30 °C). For combined hook/filament counting, EM8242 (± AnTc 0 - 4 ng/mL) and parental TH9677 were maleimide-stained, immobilized in poly-L-lysine flow cells, fixed with 4% paraformaldehyde, and filaments immunostained with anti-FliC (1:1000) followed by anti-rabbit Alexa Fluor 488 (1:1000). Images were acquired on a Nikon Eclipse Ti2 (60× Ph3 oil objective) with a 0.5 µm z-step over a 3 µm range on the 488 channel to capture all hooks in the cell. Images were pre-processed in Fiji ^48^ (channel separation, maximum-intensity projection); cells were segmented with Omnipose ^53^ and foci detected with ilastik following an established workflow ^54^, with quantification in a custom Python pipeline (github.com/SalmoLab/FliI_ATPase). Filament-positive hooks were classified by thresholding the filament-channel maximum-intensity signal within a fixed radius around each hook. Hook number per cell was used as the proxy for flagellar number throughout.

### Population motility competition assay

Fluorescently marked P*proA*-*flhDC* and P*proB*-*flhDC* strains (P*rpsM*-eGFP / P*rpsM*-mScarlet-I) were adjusted to OD600 = 1, mixed 1:1 and inoculated into soft-agar motility plates (4 h, 37 °C). Halos were imaged on a Bio-Rad ChemiDoc (Alexa 488/546), and cells sampled from four radial regions (R1-R4) were maleimide hook-stained. Strain identity was assigned from mean per-cell fluorescence (threshold 1000 in each channel; cells above or below both thresholds discarded) after phase-contrast segmentation in ilastik ^52^ and object detection in MATLAB, and hook foci centroids were assigned to their parent cells to give per-cell, per-region flagellar counts.

### Single-cell swimming competition and trajectory analysis

Pairwise swimming competitions between WT, PproA-*flhDC* and PproB-*flhDC* (six reciprocal-label combinations) were imaged in a PDMS chemotaxis chip adapted from published designs (2 mm channel, 20 µm wide, 8 µm deep; 10:1 PDMS) ^55–57^ in liquid and in 0.3% low-melting-point agarose. Trajectories were recorded at 20 frames s⁻¹ for 600 s on a Nikon Eclipse Ti2 (20× Ph2 objective, OptoSplit II dual-channel). Single-cell tracks were extracted with SwimTracker (github.com/dufourya/SwimTracker) and combined in R. Tracks were quality-filtered so that retained cells maintained a consistent speed along a smooth trajectory (|mean acceleration| < 2, speed CV < 1, mean angle < 10, mean speed 3-60 µm s^−1^, diffcoeff_cve_mean > 0.01 µm^2^ s^−1^, and 0.001 < diffcoeff_cve_runtime < 0.7 µm^2^ s^−1^, the last two excluding tracks that were either too straight or too random), and swimming vs non-swimming state was assigned from a two-component Gaussian mixture on transformed speed and effective diffusivity. Reported parameters were mean speed, effective diffusivity, swimming fraction and directional persistence (τ = 2D*eff* / v^2^), summarized as per-experiment medians for paired phenotype comparisons in agarose and liquid.

### Whole-cell proteomics

All samples were subjected to the SP3 sample preparation protocol ^58^. Ten µg of a 1:1 mixture of hydrophilic and hydrophobic carboxyl-coated paramagnetic beads (SeraMag, #24152105050250 and #44152105050250, GE Healthcare) were added for each µg of protein. Protein binding was induced by the addition of acetonitrile to a final concentration of 70% (v/v). Samples were incubated at room temperature for 10 minutes. The tubes were placed on a magnetic rack, and beads were allowed to settle for three minutes. The supernatant was discarded, and beads were rinsed thrice with 80% ethanol. Beads were resuspended in a digestion buffer containing 50 mM triethylammonium bicarbonate (Sigma, #T7408) and Trypsin and lys C (SERVA, #37283.03) in a 1:50 enzyme-to-protein ratio. Protein digestion was carried out for 14 hours at 37 °C. Afterward, the peptide supernatant was recovered and acidified with 2% ACN and 0.1% trifluoroacetic acid.Label-free DIA analyses of peptides were acquired over 120 min by an Orbitrap Exploris 480 (Thermo Scientific) coupled to a 3000 RSLC nano UPLC (Thermo Scientific) from 750 ng of peptides. Samples were loaded onto a PepMap trap cartridge (300 µm i.d. × 5 mm, C18, Thermo Scientific) with 2% acetonitrile and 0.1% TFA at a flow rate of 20 µL/min. Peptides were separated over a 25 cm analytical column (PepSep C18, 75 µm I.D., 1.5 µm). Solvent A consists of 0.1% formic acid in water. Elution was carried out at a constant flow rate of 250 nL/min for 120 min. Initially, a two-step linear gradient was applied: 5–30% solvent B (0.1% formic acid in 80% acetonitrile) over 70.5 min, then 30–45% solvent B over 13 min, followed by column washing and equilibration. The column was kept at a constant temperature of 50 °C.The MS was operated in DIA mode for single-injection quantitative measurements of individual samples with the following settings: 60k MS1 resolution, MS1 scan range 350–1250 m/z, 15k MS2 resolution, MS2 scan range 110–1600 m/z, normalised AGC target of 1000%, maximum injection time 60 ms, and fixed normalised collision energy of 30. DIA MS2 scans were performed at 12 m/z precursor isolation windows with optimized window placements from 400.4319 to 1204.7975 m/z within the precursor mass range 400–1200 m/z, with an isolation window overlap of 0.2 m/z.Raw data analysis was performed using Spectronaut (Biognosys AG, Zurich, Switzerland) version 20.1.250624.92449 in directDIA+ deep mode with reviewed UniProt databases (*Salmonella* Typhimurium strain LT2 (SGSC1412 / ATCC 700720) Proteome ID: UP000001014; 4,533 entries in UniProtKB). Methionine oxidation and Acetyl (Protein N-term) were set as a variable, and carbamidomethylation on cysteine residues was used as a static modification. The FDR for PSM-, peptide-, and protein-level was set to 0.01. All tolerances were set to dynamic for pulsar searches.

### Coarse-grained proteome-allocation model

A coarse-grained cellular-economy model was used to interpret the proteome data and predict the flagellar investment optimum. The model follows the framework of ^37^ and ^59^ and the cyanobacterial reimplementation of ^60^ (source: github.com/m-jahn/cell-economy-models), formulated as a nonlinear mixed-integer optimization in Python with the Gekko package. It represents a heterotrophic cell with eight coarse-grained protein sectors: carbon transport, carbon metabolism, electron transport chain, amino-acid biosynthesis, ribosomes/translation, lipid biosynthesis, flagella and a non-enzymatic ‘other’ sector, each with Michaelis-Menten kinetics (turnover kcat, half-saturation Km). Kinetic parameters and cellular constraints were taken from the literature and the BioNumbers ^61^ and BRENDA ^62^ databases (Supplementary Table S2). At steady state the solver maximizes growth rate by allocating ribosome capacity α to synthesize each sector i, subject to a proteome mass-fraction constraint (Σ αi = 1). Simulations varied flagellar mass fraction (0-5%), switched flagellar rotation on or off, or forced upward movement along a substrate gradient (dynamic simulation over time). The model was further constrained using data from mass spectrometry proteomics. Normalized LFQ intensities were summed up per sector and divided by total intensity to yield proteome mass fraction (R v4.5.3), which were then mapped to model sectors through a curated KEGG-pathway dictionary; the Δ*flhDC* (flagella-null) proteome was used to fine-tune kinetic parameters by iterative randomization within ±15%, minimizing the absolute log-ratio error between predicted and measured sector fractions.

### Agent-based single-cell motility simulation

To connect measured single-cell parameters to behaviour in structured media, we used a two-dimensional agent-based model implemented in Python with NumPy, pandas and Matplotlib to generate synthetic trajectories from phenotype- and medium-specific summary parameters (github.com/MPUSP/salmonella-motility-simulation). For each of 100 random seeds, we simulated 26 cells for 20 s. Motile cells underwent rotational diffusion during runs and instantaneous reorientation events with normally distributed turn angles, whereas non-motile cells underwent weak passive diffusion. The agarose-like condition contained a static field of non-overlapping circular obstacles with radii of 2.5–5.2 µm. Upon contact, cells either slid tangentially along an obstacle or entered a transient, phenotype-specific stall. The simulation was designed to illustrate, rather than predict, the experimental findings. Run speed, motile fraction and persistence time were calibrated to the measured paired-unit means. The resulting ordering of swimming speed and effective diffusivity among strains was therefore imposed by the experimental data and does not represent an independent model prediction. The model used measured summary values and did not refit parameters from raw trajectories. The reorientation-angle spread was fixed at *σ* = 1.247 rad for all six strain– medium combinations so that the mean turn magnitude matched the value of 57° measured across 8,058 turns of *E. coli* ^63^. This calibration constrains only the mean turn magnitude. It applies a three-dimensional measurement to a two-dimensional model, and the Gaussian turning distribution does not reproduce the forward-skewed shape of the measured distribution. In the agarose-like medium, the probability of stalling at each contact decreased with mean flagellar number, *N*, as *N*^−0.704^ (P*proA*, 0.210; WT, 0.177; P*proB*, 0.123), normalised to preserve their mean. The absolute probabilities and the exponent are nominal. The ordering follows the finding that additional flagella reduce trapping in agar hydrogels ^64^, which found a 1.7 ± 0.2 stall-frequency ratio per unit time in *V. alginolyticus* in 0.25% agar. The mean stall duration was fixed at the same nominal value of 0.949 s for all three strains. Reorientation was instantaneous, such that cells spent no time in a separate non-swimming reorientation state. The stall test was performed once per contact event rather than at every simulation step during an overlap. We report net displacement because contour path length did not converge with decreasing integration step. Trajectories containing a diffusive component have infinite arc length in the continuum limit, causing the summed step length to increase as the integration step decreases. When the step was reduced from 0.05 s to 0.000625 s, the simulated mean path length of WT in the agarose-like medium increased from 325 µm to 595 µm and remained unconverged, while the P*proA*/WT path-length ratio shifted from 0.54 to 1.10. By contrast, net displacement converged in both media. Across the six strain–medium combinations, the largest deviation over the same refinement series was 3.96% at 0.05 s and 1.99% at 0.0025 s. All reported simulations used an integration step of 0.0025 s and a 1,776 × 1,152 µm domain. The agarose-like medium contained 8,352 obstacle disks, corresponding to a realised obstacle area fraction of 0.187. Across 100 random seeds, the simulation reproduced 87 - 90% of the measured effective diffusivity in liquid and 69 - 76% in the agarose-like medium. The larger shortfall in agarose arose because the measured persistence time already incorporated the effects of the mesh, after which the model applied obstacles and stalls, effectively incorporating these effects twice. No such double counting occurred in liquid, where the simulation reproduced 98 - 100% of the effective diffusivity implied by its own inputs, *v*^2^*τ*/2.

### Software

Image processing used MATLAB R2022b, Fiji 2.16.0 ^48^, ImageJ 2.16.0 ^51^ and MicrobeJ 5.13p ^65^; segmentation used Omnipose 1.0.6 ^53^ and ilastik 1.4.0 ^52^; proteomics used Spectronaut 20.1.250624.92449; modelling used Python 3.14.3; Gekko 1.3.2; R 4.5.3; reference databases were NCBI, UniProt UP000001014 (4,533 entries), annotation via taxonomy ID 99287 and SalCom ^66,67^. Custom analysis code is available at the repositories cited in the relevant sections.

### Statistics and reproducibility

All inferential tests were two-sided. Statistical analyses were performed at the level of independent biological or experimental units rather than individual cells, trajectories or technical replicate measurements. The experimental unit was an independent microscopy experiment, mother-machine run, experiment day, matched soft-agar replicate or co-imaged swimming movie, as specified for each analysis. Accordingly, *n* denotes the number of experimental units. Technical replicates were averaged within each experimental unit before inference. Cell-level, trajectory-level and technical-replicate distributions were displayed to show measurement variability but were not treated as independent observations in statistical tests. Unless stated otherwise, summary values are the mean of the experimental-unit means, with ± indicating one standard deviation across units. Confidence intervals are 95% intervals. For hook and filament measurements, the mean count per cell was first calculated within each independent experiment. Conditions in the AnTc and constitutive promoter series were compared with WT using two-sided Welch’s *t*-tests on the three experiment means. Individual cells were not used as independent observations. Where several conditions within one panel were compared with the same reference, *P* values were adjusted separately within that panel using the Benjamini–Hochberg procedure and are reported as *q* values. The matched hook-versus-filament analysis was descriptive and was not tested. For batch-culture experiments, the growth rate of each culture was divided by the mean growth rate of the reference cultures measured on the same day. Reference cultures were normalized in the same manner and therefore varied around 1. Technical replicate cultures were averaged within each day, and the day mean was the unit of analysis. Conditions were compared with their matched same-day reference using two-sided paired *t*-tests across six experiment days. Benjamini–Hochberg correction was applied separately within the AnTc, constitutive promoter and mutant comparison families. Rotating and non-rotating growth penalties were compared with the matched same-day Δ*flhDC* reference using paired *t*-tests across six experiment days, followed by Benjamini-Hochberg correction. Absolute growth rates are provided in Source Data. For mother-machine experiments, individual-cell growth rates were normalized to the P*pro1*-*flhDC* mean of the same experiment for the constitutive promoter series and to the WT mean of the same experiment for the mutant series. The independent experiment, rather than the individual cell, was the experimental unit. Experiment-level summaries are provided in Source Data. For proteome-sector analyses, protein mass fractions were averaged across four biological replicates for each strain. For each non-flagellar sector, the change in sector allocation relative to Δ*flhDC* was regressed against the mean flagellar allocation of the six strains using ordinary least squares. Regression slopes are reported with model-based 95% confidence intervals. *P* values were adjusted across the seven non-flagellar sector regressions using the Benjamini-Hochberg procedure. For the AnTc soft-agar series, individual wells were normalized to the corresponding WT reference. Because WT was measured on only two of the four experiment days, this analysis was treated as descriptive and no inferential test was applied. For the constitutive promoter series, each strain was paired with the WT measurement from the same replicate and compared with WT using a two-sided paired *t*-test across six replicate pairs. The four *P* values from this panel were not adjusted for multiple comparisons. Hook-count measurements from positions within one soft-agar halo and strain-composition measurements across regions R1–R4 were obtained from single experiments and were analysed descriptively. Imaging fields quantified spatial variation within an experiment and were not considered biological replicates. For paired swimming experiments, each co-imaged movie provided one paired experimental unit because both strains were recorded simultaneously under the same conditions. Mean swimming speed, mean effective diffusivity and the fraction of time spent swimming were calculated separately for the two strains within each movie. Effective diffusivity was decomposed according to *Deff* = *v*^2^*τ*/2, where *v* is mean swimming speed and *τ* = 2*Deff*/*v*^2^ is the derived directional-persistence timescale. Ratios between paired strains were analyzed on the logarithmic scale. Point estimates are the exponentiated equal-weight means of the paired log ratios. Confidence intervals were obtained from 10,000 fixed-seed percentile bootstrap resamples of the paired movies. No separate *P* values were calculated; inference was based on the paired ratio estimates and their 95% confidence intervals. Hook counts associated with the swimming experiments were analyzed descriptively. Deterministic model outputs were not subjected to statistical testing. No inferential tests were applied to Figures 1E, 2C, 3C, 4C-F, 5, 6A, 6C, 6E or 7E-G. Because only two independent mother-machine experiments were available per strain, the mother-machine panels were analyzed descriptively. For seeded particle simulations, each seed generated 26 simulated cells per condition. Condition summaries are the medians of the 100 seed means, with intervals spanning their 2.5th-97.5th percentiles. These intervals quantify between-seed simulation variability and do not represent biological confidence intervals. Exact sample sizes, effect estimates, confidence intervals, test statistics, degrees of freedom, *P* values and *q* values are provided in the figure legends and Source Data statistics tables.

## Data and code availability

Mass-spectrometry proteomics data have been deposited in ProteomeXchange via the PRIDE ^68^ partner repository under accession PXD082813. Source data for all main and supplementary figures are provided with the paper. Processed data and figure source-data tables are archived at Zenodo under DOI 10.5281/zenodo.21952759, and the simulated trajectories of Supplementary Figure 4 under DOI 10.5281/zenodo.21953146. The analysis, simulation and figure-generation code that reproduces every panel is available at github.com/SalmoLab/salmonella-flagella-cost-benefit and archived at Zenodo under DOI 10.5281/zenodo.21951357. The coarse-grained cell-economy model is available at github.com/m-jahn/cell-economy-models, the agent-based motility simulation at github.com/MPUSP/salmonella-motility-simulation, the swimming-trajectory tracker at github.com/dufourya/SwimTracker and the hook and filament quantification pipeline at github.com/SalmoLab/FliI_ATPase.

## Acknowledgements

We thank the Erhardt, Charpentier and Dufour laboratories for continuous support and Raúl Trepel, Heidi Landmesser for expert technical assistance. This work was supported in part by a project that has received funding from the European Research Council (ERC) under the European Union’s Horizon 2020 research and innovation programme (grant agreement number 864971) and from the VolkswagenStiftung (grant number 96732). M.E. and M.J.G-Z. acknowledge funding from the Max Planck Society within the Max Planck Fellow program. The funders had no role in study design, data collection and analysis, decision to publish, or preparation of the manuscript.

## Author contributions

M.J.G-Z., Y.S.D. and M.E. conceived the project and designed the study; M.J.G-Z. and M.E. wrote the paper; all authors contributed to revisions; M.J.G-Z., M.J., K.A., F.K., J.L.F. and S.D. performed the experiments; M.J.G-Z., M.J., K.A., J.L.F., S.D., P.F.P., Y.S.D. and M.E. analyzed and interpreted the data; K.T.H., E.C., Y.S.D. and M.E. contributed funding and resources.

