## Supplementary Information for "The cost-benefit trade-off of peritrichous flagellation in bacteria"

**Supplementary Information Table of Contents**

Supplementary Information Figure S1 - S6
Supplementary Information Table S1 - S4
Supplementary Information References

#### Supplementary Information Figures

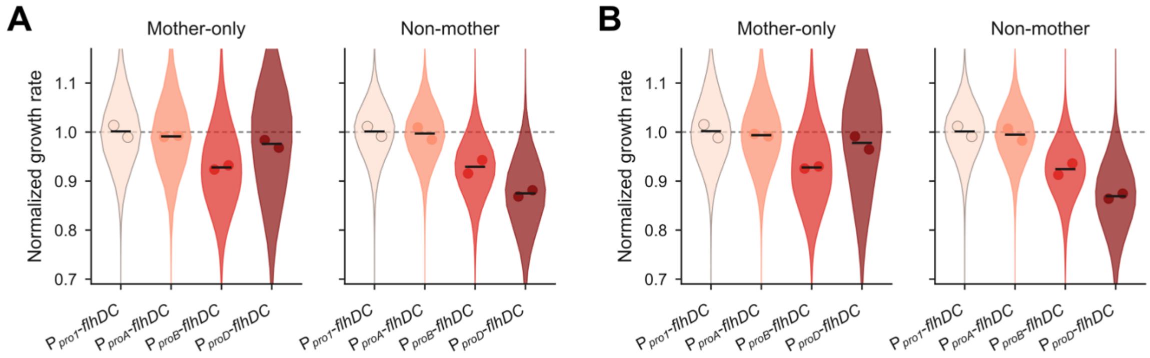


**Supplementary Figure S1. Single-cell growth across two analysis windows.** Mother-machine growth rates separated by lineage and analysis window. Mother cells occupied the closed ends of channels; other cells were their descendants. A, Analysis from 200 - 800 min, comprising 126,934 division cycles. B, Analysis from 180 - 480 min, comprising 65,967 division cycles. Violins show pooled growth rate distributions; points are the means of two independent experiments and bars are their means.

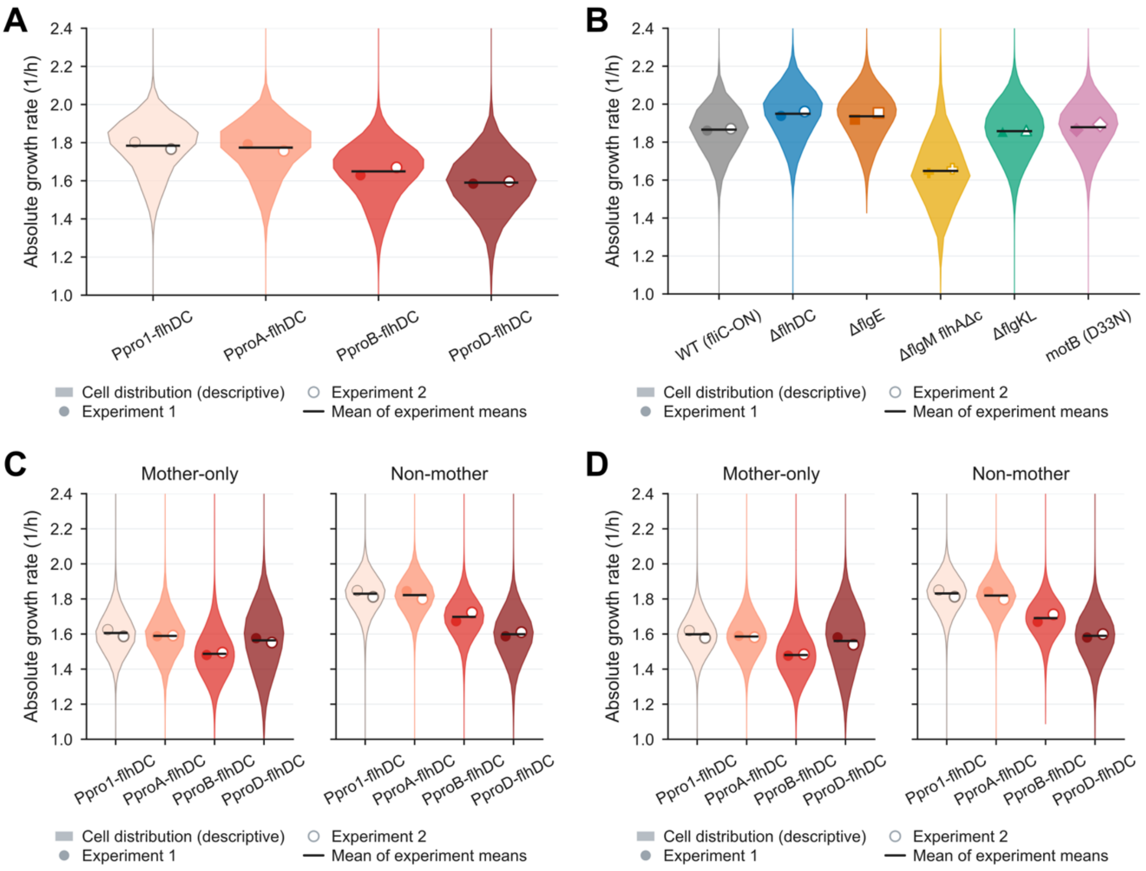


**Supplementary Figure S2. Absolute single-cell growth rates.** Mother-machine growth rates in absolute units, for every dataset shown in Figures 2C and 3C and Supplementary Figure S1. A, Promoter series, all cells, 200 - 800 min, comprising 126,934 division cycles. B, Flagellar assembly mutants, all cells, 200 - 800 min, comprising 110,983 division cycles. Promoter series separated into mother and non-mother lineages, over 200 - 800 min (C) and 180 - 480 min (D). C comprises 27,487 mother and 99,447 non-mother cycles, and D comprises 14,205 mother and 51,762 non-mother cycles. Violins show the pooled growth rate distribution. The filled and the open mark are the means of the two independent mother-machine experiments of that strain, and the bar is the mean of the two experiment means. Per-experiment means, standard deviations and cycle counts are given in Supplementary Table S1.

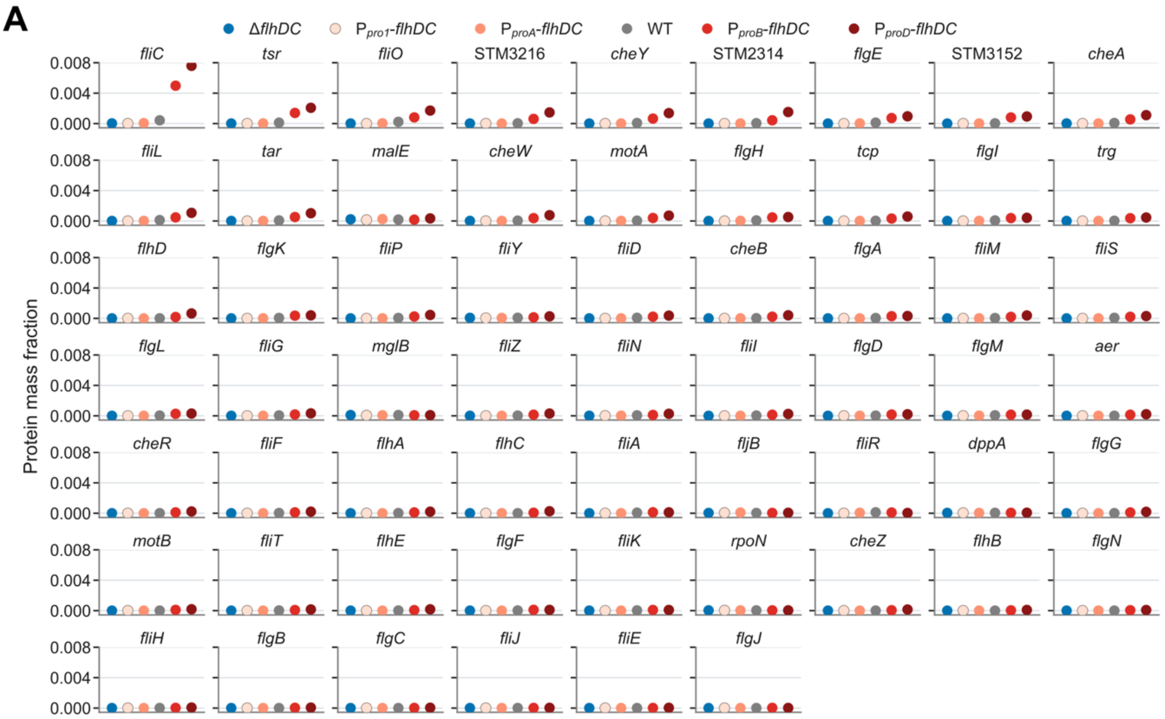


**Supplementary Figure S3. Protein-level composition of the flagellar sector.** Mass fraction of each flagellar and chemotaxis protein in Δ*flhDC*, $P_{pro1}$-*flhDC*, $P_{proA}$-*flhDC*, WT, $P_{proB}$-*flhDC* and $P_{proD}$-*flhDC*. Values are means of four biological replicates. Sector assignments are as in Figure 4.

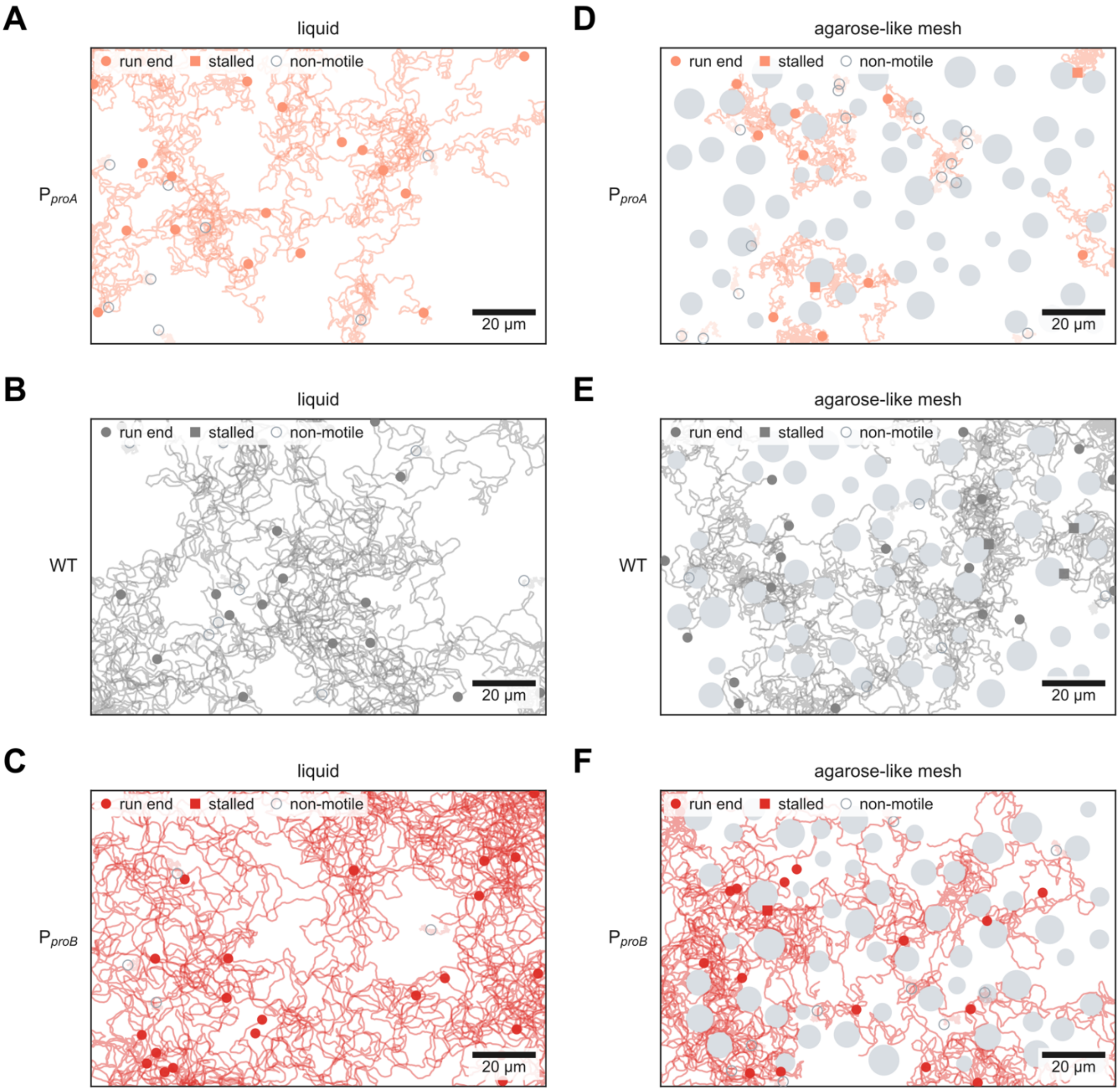


**Supplementary Figure S4. Representative simulated trajectories.** Representative trajectories from one seed of the particle simulation in Figure 5D, E. P$_{proA}$-*flhDC*, WT and P$_{proB}$-*flhDC* are shown in liquid and in the agarose-like obstacle mesh. Grey discs are obstacles. Labels give the input swimming speed, effective diffusivity and motile fraction. The figure is illustrative and carries no statistical test.

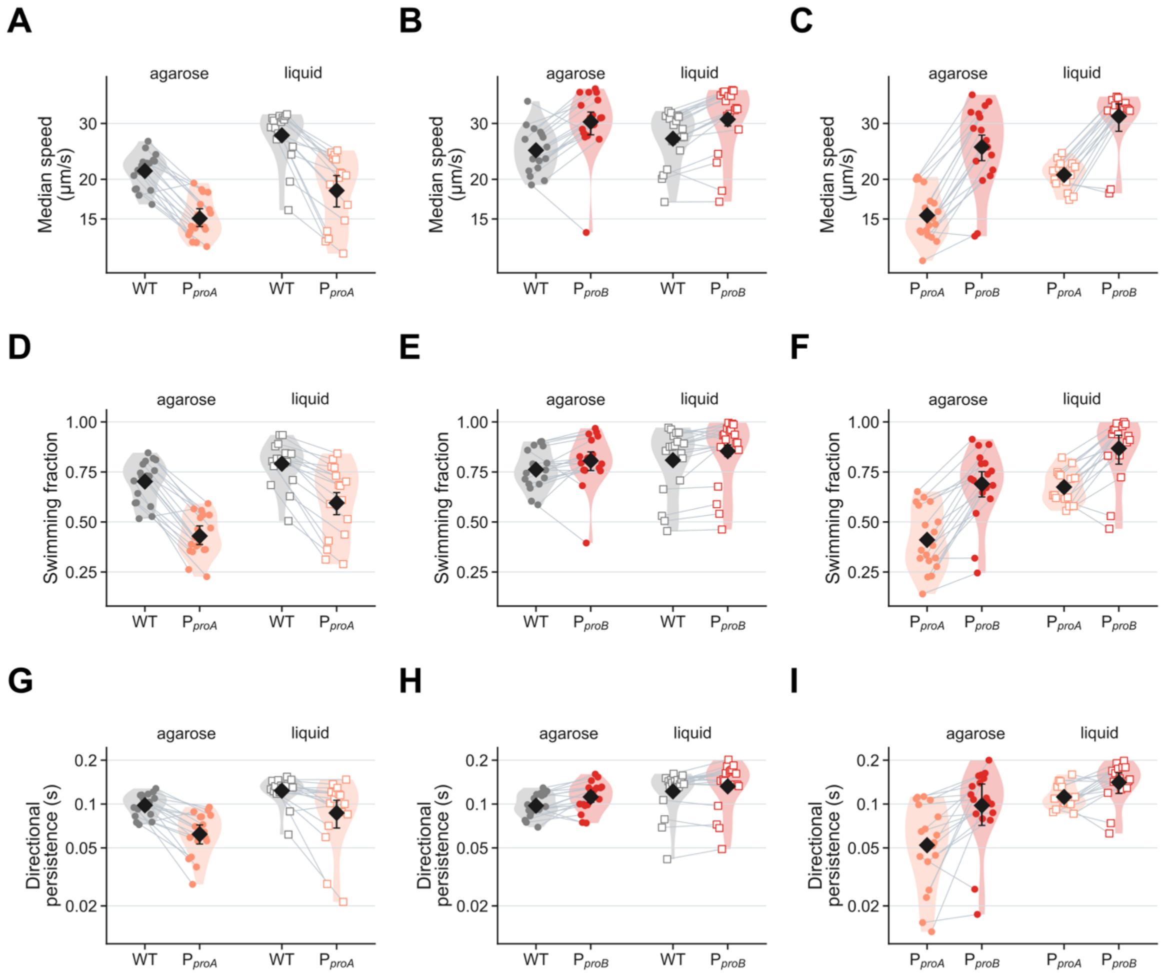


**Supplementary Figure S5. Paired swimming measurements by phenotype.** Swimming measurements from the movies analyzed in Figure 7. A-C, Mean swimming speed. D-F, Mean effective diffusivity. G-I, Fraction of time spent swimming. Columns show WT against $P_{proA}$-*flhDC*, WT against $P_{proB}$-*flhDC*, and $P_{proA}$-*flhDC* against $P_{proB}$-*flhDC*. Symbols represent movies and lines connect co-imaged strains. Filled circles, agarose; open squares, liquid. Paired estimates and confidence intervals are shown in Figure 7D.

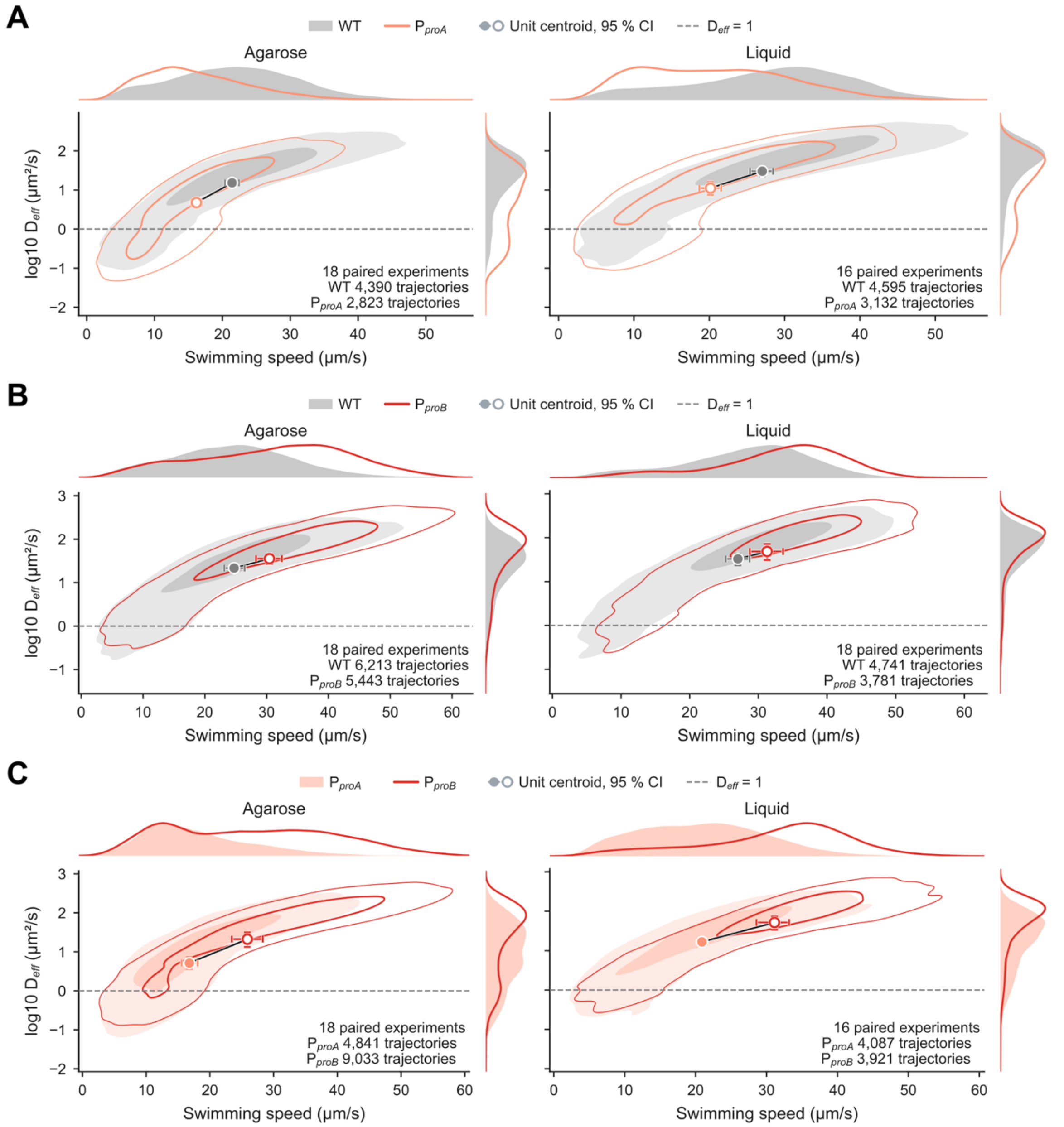


**Supplementary Figure S6. Speed and effective-diffusivity distributions.** Pooled single-cell swimming speed and effective diffusivity for the three strain pairs analyzed in Figure 7. A-C, WT against $P_{proA}$-*flhDC*, WT against $P_{proB}$-*flhDC*, and $P_{proA}$-*flhDC* against $P_{proB}$-*flhDC*. Swimming and non-swimming cells are shown separately; contours indicate cell density. Paired inference is shown in Figure 7A-D

### Supplementary Information Tables

**Supplementary Table S1. Absolute single-cell growth rates.** Mean growth rate of each independent mother-machine experiment, their mean, and the pooled number of division cycles, for the promoter series and the flagellar assembly mutants. One value is one completed birth-to-division cycle: a lineage contributes many cycles. Standard deviations across cycles are provided with Source Data.

| Strain | Lineage | Window (min) | Exp. 1 (1/h) | Exp. 2 (1/h) | Mean (1/h) | Cycles |
|---|---|---|---|---|---|---|
| Ppro1-flhDC | All cells | 180-480 | 1.803 | 1.765 | 1.784 | 19,787 |
| PproA-flhDC | All cells | 180-480 | 1.789 | 1.753 | 1.771 | 17,831 |
| PproB-flhDC | All cells | 180-480 | 1.626 | 1.661 | 1.643 | 13,462 |
| PproD-flhDC | All cells | 180-480 | 1.581 | 1.585 | 1.583 | 14,887 |
| Ppro1-flhDC | Mother-only | 180-480 | 1.620 | 1.578 | 1.599 | 4,025 |
| PproA-flhDC | Mother-only | 180-480 | 1.590 | 1.582 | 1.586 | 3,660 |
| PproB-flhDC | Mother-only | 180-480 | 1.476 | 1.484 | 1.480 | 3,034 |
| PproD-flhDC | Mother-only | 180-480 | 1.582 | 1.540 | 1.561 | 3,486 |
| Ppro1-flhDC | Non-mother | 180-480 | 1.851 | 1.813 | 1.832 | 15,762 |
| PproA-flhDC | Non-mother | 180-480 | 1.842 | 1.797 | 1.819 | 14,171 |
| PproB-flhDC | Non-mother | 180-480 | 1.670 | 1.712 | 1.691 | 10,428 |
| PproD-flhDC | Non-mother | 180-480 | 1.580 | 1.599 | 1.590 | 11,401 |
| Ppro1-flhDC | All cells | 200-800 | 1.802 | 1.766 | 1.784 | 38,650 |
| PproA-flhDC | All cells | 200-800 | 1.792 | 1.757 | 1.774 | 34,899 |
| PproB-flhDC | All cells | 200-800 | 1.629 | 1.670 | 1.649 | 25,672 |
| PproD-flhDC | All cells | 200-800 | 1.585 | 1.596 | 1.590 | 27,713 |
| Ppro1-flhDC | Mother-only | 200-800 | 1.625 | 1.587 | 1.606 | 7,883 |
| PproA-flhDC | Mother-only | 200-800 | 1.588 | 1.591 | 1.589 | 7,167 |
| PproB-flhDC | Mother-only | 200-800 | 1.480 | 1.494 | 1.487 | 5,912 |
| PproD-flhDC | Mother-only | 200-800 | 1.576 | 1.552 | 1.564 | 6,525 |
| Ppro1-flhDC | Non-mother | 200-800 | 1.848 | 1.812 | 1.830 | 30,767 |
| PproA-flhDC | Non-mother | 200-800 | 1.845 | 1.799 | 1.822 | 27,732 |
| PproB-flhDC | Non-mother | 200-800 | 1.673 | 1.723 | 1.698 | 19,760 |
| PproD-flhDC | Non-mother | 200-800 | 1.587 | 1.611 | 1.599 | 21,188 |
| WT (fliC-ON) | All cells | 200-800 | 1.861 | 1.871 | 1.866 | 16,990 |
| ΔflhDC | All cells | 200-800 | 1.938 | 1.961 | 1.950 | 20,543 |
| ΔflgE | All cells | 200-800 | 1.919 | 1.954 | 1.936 | 19,593 |
| ΔflgM flhAΔc | All cells | 200-800 | 1.637 | 1.660 | 1.648 | 15,600 |
| ΔflgKL | All cells | 200-800 | 1.854 | 1.861 | 1.858 | 19,162 |
| motB (D33N) | All cells | 200-800 | 1.864 | 1.894 | 1.879 | 19,095 |
| WT (fliC-ON) | Mother-only | 200-800 | 1.777 | 1.772 | 1.774 | 3,354 |
| ΔflhDC | Mother-only | 200-800 | 1.806 | 1.820 | 1.813 | 3,888 |
| ΔflgE | Mother-only | 200-800 | 1.778 | 1.823 | 1.801 | 3,950 |
| ΔflgM flhAΔc | Mother-only | 200-800 | 1.600 | 1.682 | 1.641 | 3,460 |
| ΔflgKL | Mother-only | 200-800 | 1.759 | 1.763 | 1.761 | 3,841 |
| motB (D33N) | Mother-only | 200-800 | 1.775 | 1.777 | 1.776 | 3,869 |
| WT (fliC-ON) | Non-mother | 200-800 | 1.881 | 1.895 | 1.888 | 13,636 |
| ΔflhDC | Non-mother | 200-800 | 1.968 | 1.995 | 1.981 | 16,655 |
| ΔflgE | Non-mother | 200-800 | 1.954 | 1.987 | 1.970 | 15,643 |
| ΔflgM flhAΔc | Non-mother | 200-800 | 1.648 | 1.653 | 1.651 | 12,140 |
| ΔflgKL | Non-mother | 200-800 | 1.877 | 1.887 | 1.882 | 15,321 |
| motB (D33N) | Non-mother | 200-800 | 1.886 | 1.924 | 1.905 | 15,226 |

**Supplementary Table S2. Parameters of the eight-sector proteome-allocation model.** Values are those used in the simulations. Sectors are carbon transport (Tra), carbon metabolism (Cbn), electron transport chain (Etc), amino-acid biosynthesis (Aab), ribosomes and translation (Rib), lipid biosynthesis (Lpb), flagella (Fla) and a non-enzymatic other sector (Oth). Sector protein amounts were derived from the measured proteome mass fractions and are given in Figure 4D. Cell length and radius were constrained to the stated intervals; cell volume and surface area were calculated from them for a cylindrical cell.

| Component | Parameter | Value | Unit | Reference |
|---|---|---|---|---|
| ***Cell and environment*** | | | | |
| cell | protein density | $8 \times 10^9$ | aa $\mu m^{-3}$ | [1] |
| cell | length | 2–5 | µm | [2] |
| cell | radius | 0.25-0.75 | µm | [2] |
| cell | volume | calculated from length and radius | $\mu m^3$ | — |
| cell | surface area | calculated from length and radius | $\mu m^2$ | — |
| cell | flagellar rotation frequency | 100 | $s^{-1}$ | [3] |
| cell | flagellar energy cost | 250 | ATP per rotation | [4]; [5]; [6] |
| cell | maximum swimming speed | 35 | µm $s^{-1}$ | BioNumbers 106818; [7] |
| cell | half-saturation of the swimming response | 10 | model units of c(Fla) | assumed |
| cell | run-and-tumble drift efficiency | 0.01 | dimensionless | [8] |
| cell | glucose diffusion coefficient | 600 | $\mu m^2$ $s^{-1}$ | BioNumbers 104089 |
| cell | initial glucose concentration | 5 | mM | assumed |
| cell | glucose profile | $C(x,t) = C_0 \cdot \mathrm{erfc}(x / (2\sqrt{(D \cdot t)}))$ | mM | Crank J. The Mathematics of Diffusion, 2nd ed. Oxford University Press, 1975 |
| ***Sector turnover, k_cat*** | | | | |
| Tra | k_cat (a) | 219 | molec $s^{-1}$ | BioNumbers 103693; 114686 |
| Cbn | k_cat (a) | 105 | molec $s^{-1}$ | BioNumbers 100996; 100997; 101029 |
| Etc | k_cat (a) | 20 | molec $s^{-1}$ | BioNumbers 111975 |
| Aab | k_cat (a) | 135 | molec $s^{-1}$ | BRENDA EC 6.3.1.2 / 2.6.1.1 / 1.4.1.1 / 1.1.1.3 / 1.4.1.13 / 2.1.2.1 |
| Rib | k_cat (a) | 7.09 | aa $s^{-1}$ | BioNumbers 100059; [1] |
| Lpb | k_cat (a) | 5.69 | molec $s^{-1}$ | BRENDA EC 6.4.1.2 |
| Fla | k_cat | 25,000 | ATP $s^{-1}$ | derived, see (b) |
| ***Sector half-saturation, K_m*** | | | | |
| Tra | K_m (a) | 2.3 | mM | BioNumbers 110956 |
| Cbn | K_m (a) | 0.02 | mM | BioNumbers 104956; 104957 |
| Etc | K_m (a) | 0.1 | mM | BioNumbers 104891 |
| Aab | K_m (a) | 0.32 | mM | BRENDA EC 6.3.1.2 / 2.6.1.1 / 1.4.1.1 / 1.1.1.3 / 1.4.1.13 / 2.1.2.1 |
| Rib | K_m (a) | 1.57 | mM | assumed |
| Lpb | K_m (a) | 0.31 | mM | BRENDA EC 2.1.3.15; EC 6.4.1.2 |
| Fla | K_m | 1 | mM | assumed |
| ***Sector Hill coefficient*** | | | | |
| all sectors | Hill coefficient | 1 | dimensionless | assumed |
| ***Sector protein size*** | | | | |
| Tra | protein size | 1,000 | aa | UniProt; PDB |
| Cbn | protein size | 10,000 | aa | UniProt |
| Etc | protein size | 10,000 | aa | UniProt |
| Aab | protein size | 5,000 | aa | UniProt |

| | | | | |
|---|---|---|---|---|
| Rib | protein size | 7,536 | aa | BioNumbers 110218 |
| Lpb | protein size | 2,000 | aa | UniProt |
| Fla | protein size | 7,200,000 | aa | [9]; [10]; see (c), (d) |
| Oth | protein size | 350 | aa | assumed |
| ***Specific membrane area, spA*** | | | | |
| Tra | spA | $5 \times 10^{-5}$ | $\mu m^2$ | [11]; PDB 9HNP |
| Fla | spA | $5 \times 10^{-4}$ | $\mu m^2$ | [10]; PDB 7NVG |
| Etc | spA | $5 \times 10^{-3}$ | $\mu m^2$ | assumed |
| membrane lipid | spA | $1 \times 10^{-5}$ | $\mu m^2$ | assumed |

**(a)** k_cat and K_m were optimized against the ΔflhDC proteome by iterative randomization within ±15–20% of their starting values, minimizing the absolute log-ratio error between predicted and measured sector mass fractions. k_cat(Fla) was held fixed.

**(b)** k_cat(Fla) = 250 ATP per rotation × 100 rotations $s^{-1}$. The energy cost follows from 572 to 1,100 protons per rotation at 3.33 protons per ATP.

**(c)** Fla protein size = 200,000 aa for the basal body, hook, rotor and stator, plus 2,000 flagellin subunits per μm over a 7 μm filament at 500 aa per subunit.

**(d)** Calculation for the wild-type flagellar investment as quoted in the Discussion. Whole-cell proteomics measures the intracellular pool, so the external filament is absent from the measured flagellar sector. Mass balance per cell: FliC is 51,611 Da (UniProt P06179), or $8.57 \times 10^{-5}$ fg per subunit; at 2,000 subunits $\mu m^{-1}$ a 7 μm filament carries 14,000 subunits and 1.20 fg of flagellin; three to four filaments per cell therefore carry 3.60 to 4.80 fg, or 1.80% to 2.40% of the 200 fg of total cell protein [12]. Retained non-filament flagellar protein is the measured sector minus measured FliC, 0.351% − 0.043% = 0.308%. Corrected investment = 2.11% to 2.71%. Measured FliC is 0.043% against the 1.80% to 2.40% expected, so 98% of filament flagellin is lost before analysis. The estimate scales with filament length and with total cell protein: at 5 μm and 10 μm it becomes 1.59-2.02% and 2.88-3.74%, and at 150 fg and 260 fg of cell protein 2.71-3.51% and 1.69-2.15%.

**Supplementary Table S3. Bacterial strains used in this study.**

All strains are *Salmonella enterica* serovar Typhimurium LT2 derivatives.

| Strain | Relevant genotype | Promoter / phenotype | Source |
|---|---|---|---|
| **TH437** | LT2 wild type | parental wild type | J. Roth |
| **TH5861** | Δ*hin*-5717::FCF (FliC-ON) | WT reference, Figure 3 | Lab collection |
| **TH9677** | Δ*hin*-5717::FRT (FliC-ON) *flgE*6506(S171C) | WT reference, Figures 1, 2, 6A, 6C | Lab collection |
| **EM4208** | Δ*flgE*1204 Δ*hin*-5717::FCF (FliC-ON) | basal body, no hook | Lab collection |
| **EM4209** | Δ*flgKL* Δ*hin*-5717::FCF (FliC-ON) | flagellin secreted, no filament | Lab collection |
| **EM8242** | Δ*hin*-5717::FRT (FliC-ON) *flgE*6506(S171C) P*flhDC*5451::Tn10dTc[del-25] Δ*tetA*4 (ΔAA6–396) | P*tetA-flhDC* (AnTc-inducible) | Lab collection |
| **EM8325** | Δ*hin*-5717::FRT (FliC-ON) *flgE*6506(S171C) Δ*rflP*::FRT | WT, Δ*rflP* | Lab collection |
| **EM8513** | Δ*hin*-5717::FRT (FliC-ON) *flgE*6506(S171C) P*flhDC*23263::PproD-RBS (Δbp −598 to AUG of *flhD*) | P*proD-flhDC* | This study |
| **EM9660** | Δ*hin*-5717::FRT (FliC-ON) *flgE*6506(S171C) P*flhDC*23253::PproB-RBS (Δbp −598 to AUG of *flhD*) | P*proB-flhDC* | Lab collection |
| **EM9661** | Δ*hin*-5717::FRT (FliC-ON) *flgE*6506(S171C) P*flhDC*23254::PproA-RBS (Δbp −598 to AUG of *flhD*) | P*proA-flhDC* | Lab collection |
| **EM9662** | Δ*hin*-5717::FRT (FliC-ON) *flgE*6506(S171C) P*flhDC*23255::Ppro1-RBS (Δbp −598 to AUG of *flhD*) | P*pro1-flhDC* | Lab collection |
| **EM14795** | Δ*hin*-5717::FRT (FliC-ON) *flgE*6506(S171C) P*flhDC*23253::PproB-RBS (Δbp −598 to AUG of *flhD*) Δ*rflP*::FRT | P*proB-flhDC*, Δ*rflP* | This study |
| **EM14796** | Δ*hin*-5717::FRT (FliC-ON) *flgE*6506(S171C) P*flhDC*23254::PproA-RBS (Δbp −598 to AUG of *flhD*) Δ*rflP*::FRT | P*proA-flhDC*, Δ*rflP* | This study |
| **EM15549** | Δ*hin*-5717::FRT (FliC-ON) *flgE*6506(S171C) P*flhDC*23255::Ppro1-RBS (Δbp −598 to AUG of *flhD*) Δ*rflP*::FRT | P*pro1-flhDC*, Δ*rflP* | This study |
| **EM15550** | Δ*hin*-5717::FRT (FliC-ON) *flgE*6506(S171C) P*flhDC*23263::PproD-RBS (Δbp −598 to AUG of *flhD*) Δ*rflP*::FRT | P*proD-flhDC*, Δ*rflP* | This study |
| **EM16106** | Δ*hin*-5717::FRT (FliC-ON) *flgE*6506(S171C) Δ*rflP*::FRT Δ*amyA*::P*rpsM*-eGFP | WT, Δ*rflP*, eGFP | This study |
| **EM16107** | Δ*hin*-5717::FRT (FliC-ON) *flgE*6506(S171C) Δ*rflP*::FRT Δ*amyA*::P*rpsM*-mScarlet-I | WT, Δ*rflP*, mScarlet-I | This study |
| **EM16114** | Δ*hin*-5717::FRT (FliC-ON) *flgE*6506(S171C) P*flhDC*23254::PproA-RBS (Δbp −598 to AUG of *flhD*) Δ*rflP*::FRT Δ*amyA*::P*rpsM*-eGFP | P*proA-flhDC*, Δ*rflP*, eGFP | This study |
| **EM16115** | Δ*hin*-5717::FRT (FliC-ON) *flgE*6506(S171C) P*flhDC*23254::PproA-RBS (Δbp −598 to AUG of *flhD*) | P*proA-flhDC*, Δ*rflP*, mScarlet-I | This study |

| | | | |
|---|---|---|---|
| | Δ*rflP*::FRT Δ*amyA*::P*rpsM*-mScarlet-I | | |
| **EM16309** | Δ*hin*-5717::FRT (FliC-ON) *flgE*6506(S171C) P*flhDC*23253::PproB-RBS (Δbp –598 to AUG of *flhD*) Δ*rflP*::FRT Δ*amyA*::P*rpsM*-eGFP | P*proB*-*flhDC*, Δ*rflP*, eGFP | This study |
| **EM16310** | Δ*hin*-5717::FRT (FliC-ON) *flgE*6506(S171C) P*flhDC*23253::PproB-RBS (Δbp –598 to AUG of *flhD*) Δ*rflP*::FRT Δ*amyA*::P*rpsM*-mScarlet-I | P*proB*-*flhDC*, Δ*rflP*, mScarlet-I | This study |
| **EM16223** | Δ*flhDC*7884::FRT Δ*hin*-5717::FCF (FliC-ON) | Δ*flhDC* (non-flagellated) | This study |
| **EM16224** | *motB*(D33N) Δ*hin*-5717::FCF (FliC-ON) | non-rotating motor | This study |
| **EM16587** | Δ*flgM*5628::FRT Δ*hin*-5717::FCF (FliC-ON) Δ*flhA*23039 (FlhA_C deletion, RBS FlhE duplicated) | flagellin made, not secreted | This study |

**Supplementary Table S4. Strains used for each figure panel.**

| Figure | Panel | Strains | Phenotype |
|---|---|---|---|
| Figure 1 | A-E | EM8242; TH9677 | P*tetA*-*flhDC* (0–4 ng $ml^{-1}$ AnTc); WT |
| Figure 1 | F-G | EM9662; EM9661; EM9660; EM8513; TH9677 | Ppro1; PproA; PproB; PproD; WT |
| Figure 2 | A | EM8242 (six AnTc concentrations); TH9677 | P*tetA*-*flhDC*; WT |
| Figure 2 | B | EM9662; EM9661; EM9660; EM8513; TH9677 | Ppro1; PproA; PproB; PproD; WT |
| Figure 2 | C | EM9662; EM9661; EM9660; EM8513 | promoter series, normalised to Ppro1 |
| Figure 3 | B, C | EM16223; EM4208; EM16587; EM4209; EM16224; TH5861 | Δ*flhDC*; Δ*flgE*; Δ*flgM flhA*Δc; Δ*flgKL*; *motB*(D33N); WT |
| Figure 3 | E | EM16224; TH5861; model at 5% allocation | *motB*(D33N); WT |
| Figure 4 | B-F | EM9662; EM9661; EM9660; EM8513; TH9677; EM16223 | Ppro1; PproA; PproB; PproD; WT; Δ*flhDC* |
| Figure 6 | A | EM8242; TH9677 | P*tetA*-*flhDC*; WT |
| Figure 6 | B | EM15549; EM14796; EM14795; EM15550; EM8325 | Ppro1; PproA; PproB; PproD; WT |
| Figure 6 | C | TH9677 | WT halo, three radial positions |
| Figure 6 | D, E | EM16115; EM16309 | PproA-mScarlet-I; PproB-eGFP |
| Figure 7 | A, E | EM16106/EM16107 vs EM16114/EM16115 | WT vs PproA, reciprocal labels |
| Figure 7 | B, F | EM16106/EM16107 vs EM16309/EM16310 | WT vs PproB, reciprocal labels |
| Figure 7 | C, G | EM16114/EM16115 vs EM16309/EM16310 | PproA vs PproB, reciprocal labels |
| Supplementary Figure S1 | A, B | EM9662; EM9661; EM9660; EM8513 | promoter series, two analysis windows |
| Supplementary Figure S2 | A, C, D | EM9662; EM9661; EM9660; EM8513 | promoter series, absolute rates |
| Supplementary Figure S2 | B | EM16223; EM4208; EM16587; EM4209; EM16224; TH5861 | assembly mutants, absolute rates |
| Supplementary Figure S3 | A | as Figure 4 | promoter series, WT and Δ*flhDC* |
| Supplementary Figure S5 | A-I | as Figure 7 | WT; PproA; PproB |
| Supplementary Figure S6 | A-C | as Figure 7 | WT; PproA; PproB |

#### Supplementary Information References
